# A step in the deep evolution of Alvinellidae (Annelida: Polychaeta): A phylogenomic comparative approach based on transcriptomes

**DOI:** 10.1101/2023.07.24.550320

**Authors:** Pierre-Guillaume Brun, Stéphane Hourdez, Marion Ballenghien, Yadong Zhou, Jean Mary, Didier Jollivet

**Author notes:** JM and DJ are joint senior authors.

## Abstract

The Alvinellidae are a family of worms that are endemic to deep-sea hydrothermal vents in the Pacific and Indian Oceans. These annelid worms, a sister group to the Ampharetidae, occupy a wide range of thermal habitats. The family includes the most thermotolerant marine animals described to date such as the Pompeii worm *Alvinella pompejana*, and other species living at much lower temperatures such as *Paralvinella grasslei* or *Paralvinella pandorae*. The phylogeny of this family has not been studied extensively. It is, however, a complex case where molecular phylogenies have given conflicting results, especially concerning the monophyletic or polyphyletic nature of the genus *Paralvinella*.

We carried out a comprehensive study of the phylogeny of this family using the best molecular data currently available from RNAseq datasets, leading to several hundred orthologous transcripts for 11 of the 14 species currently described or in description. The results obtained by the two most popular phylogenetic inference models (using either gene concatenation with maximum likelihood, or a coalescent-based model from gene trees) were compared using a series of ampharetid and terebellid outgroups.

Our study shows that the global phylogenetic signal favors the hypothesis of paraphyly for the *Paralvinella* genus, with *P. pandorae* being sister species of the other Alvinellidae. However, a high number of gene trees also supports the hypothesis of alternative trees in which the monophyly of the *Paralvinella* genus, as initially proposed by Desbruyères and Laubier, is valid with the species *P. pandorae* and *Paralvinella unidentata* being sister species. According to molecular dating, the radiation of the Alvinellidae was rapid and took place in a short period of time between 80 and 110 million years ago. This is reflected at the genomic scale by gene trees equally separated between different phylogenetic hypothesis, showing high rates of incomplete lineage sorting between the first lineages of the Alvinellidae and probable gene transfers. Although different genomic regions seem to have different phylogenetic stories in the early step of the alvinellid radiation, our study supports the view that the two *P. pandorae* species can be grouped into a separate genus (possibly *Nautalvinella*) and that the *Miralvinella* subgenus, defined by Desbruyères and Laubier, is not valid anymore.

## 1. Introduction

After the discovery of hydrothermal vents and their associated communities on the Galapagos rift in the late 1970s, the Pompeii worm *Alvinella pompejana* Desbruyères and Laubier (1980) was one of the first emblematic species collected from the walls of vent chimneys following the sampling of a ”black smoker” at 21°N on the East Pacific Rise (EPR) (Monaco and Prouzet, 2015). Two morphological types living in close association, initially viewed as ontogenic forms, were later found to represent two distinct species, *Alvinella pompejana* and *A. caudata* Desbruyères and Laubier (1986) living in syntopy (Jollivet and Hourdez, 2020). While the two *Alvinella* species were initially described as aberrant forms in the family Ampharetidae, some obvious specificities, such as the lack of separation between the head and the rest of the body, a process of cephalization of the gills, uncini with a reduced number of teeth and notopodia with simple dorsal chaetae, led the authors to suggest the creation of the subfamily initially named Alvinellinae (Desbruyères and Laubier, 1980, 1982). Following the discovery of the species *Paralvinella grasslei* Desbruyères and Laubier (1982), at several sites in the EPR, the Galapagos rift and the Guaymas basin (eastern Pacific), the Alvinellinae were split into two genera, *Alvinella* and *Paralvinella*. The distinction between the two genera was based in particular on the shape of the mouth apparatus and the shape of the secondary filaments of the gills, which are lamellar in *Alvinella* and filamentous in *Paralvinella* Desbruyères and Laubier (1982).

Subsequently, new species of Alvinellinae were discovered, all endemic to hydrothermal vents in the Pacific Ocean. In the south-east Pacific, two additional species were found and described as *Paralvinella pandorae irlandei* Desbruyères and Laubier (1986) (EPR) and *Paralvinella bactericola* Desbruyères and Laubier (1991) (Guaymas basin). In the north Pacific, several new species were described from the Juan de Fuca Ridge with *Paralvinella pandorae pandorae* Desbruyères and Laubier (1986), *Paralvinella palmiformis* Desbruyères and Laubier (1986), *Paralvinella dela* Detinova (1988), as well as *Paralvinella sulfincola* Desbruyères and Laubier (1993) found on top of vent chimneys. Finally, three additional species were added to the family from expeditions in the western Pacific: *Paralvinella hessleri* Desbruyères and Laubier (1989) found in the Marianas back-arc basins and Okinawa Trough, and *Paralvinella fijiensis* Desbruyères and Laubier (1993) and *Paralvinella unidentata* Desbruyères and Laubier (1993) found in the south-west Pacific (North-Fiji and Lau basins).

As the number of species in the subfamily increased, subsequent studies using both morphological characters and molecular data demonstrated that inside Terebelliformia, alvinellid worms form a monophyletic group, sister to the Ampharetidae but not nested within. The family was accordingly renamed Alvinellidae (Rousset et al., 2003; Glasby et al., 2004; Stiller et al., 2020). Recently, a new species, *Paralvinella mira* Han, Zhang, Wang & Zhou, 2021, genetically close to *P. hessleri*, has been described outside of the Pacific Ocean, from the hydrothermal vents of the Carlsberg Ridge in the Indian Ocean (Han et al., 2021). The Alvinellid worms’ distribution has thus expanded beyond Pacific waters, supporting the purported observation of a small population of alvinellid worms in the Solitaire field on the Central Indian Ridge, although they have been neither sampled nor taxonomically resolved (Nakamura et al., 2012). Finally, another species, whose description is still in progress, was first discovered in 2011 from the Nifonea vent site along the volcanic arc of Vanuatu but also in the Manus Basin. This species is referred to as *Paralvinella* sp. nov. in this study.

Alvinellid worms probably have a long evolutionary history of speciation, which could date back between 70 and 200 millions of years ago (Jollivet and Hourdez, 2020). The family has diversified to cope with a wide variety of vent conditions. Alvinellidae notably include two of the most thermotolerant animals known to date, namely *A. pompejana* and its ecological homolog *P. sulfincola* thriving between 40 and 50°C (Girguis and Lee, 2006; Ravaux et al., 2013), while species such as *P. grasslei* are comfortable in temperatures around 15°C (Cottin et al., 2008) or lesser.

The ecological diversity of these closely-related worms is particularly intriguing. The question of the evolutionary path taken by the alvinellids naturally arises, as well as the different stages that allowed its dispersal throughout the Pacific to the Indian Ocean, including their putative initial link with the hydrothermal paleocoastal environment. A necessary step in the understanding of the evolutionary history of alvinellid species and their relationships during the early steps of the family radiation is the establishment of a robust phylogeny of the Alvinellidae, including all currently known and described species. Several phylogenies of the Alvinellidae have been already proposed using a variety of characters (morphological and ecological homologies, Nei’s genetic divergence of allozymes, or molecular phylogenies using orthologous transcripts). Based on detailed observations of morphological traits, Desbruyères and Laubier divided the family into two monophyletic genera, *Alvinella* and *Paralvinella* (Desbruyères and Laubier, 1993). Species of the genus *Alvinella* have gills with flattened-leaf secondary filaments, the anterior fourth and fifth setigers are modified with the insertion of two pairs of hooks, and a pair of thick tentacles is inserted on the buccal apparatus in males. Species of the genus *Paralvinella* have stalked bipennate gills with two rows of cylindrical filaments and no filaments at the tip of the stem, the seventh setiger is modified with a single pair of hooks, and the modified buccal tentacles of males are different (either three-lobed, coiled and tapered, or absent) (Jollivet and Hourdez, 2020). Other non-morphological differences between the two genera have been noted: *Alvinella* species secrete thick parchment-like tubes including inorganic material and have epibiotic bacteria, whereas *Paralvinella* species secrete mucus tubes or cocoons (with the exception of *P.* sp. nov., personal communication) and are devoid of filamentous epibiotic bacteria (Desbruyères and Laubier, 1991).

Within the *Paralvinella* species, Desbruyères and Laubier further proposed to group the species *P. p. pandorae*, *P. p. irlandei* and *P. unidentata* within the subgenus *Nautalvinella*, considering the comb-like insertion of the secondary filaments of the gills as well as the lack of modified tentacles on the buccal apparatus of the males. However, some peculiarities are noted in the species *P. unidentata*, notably the leaf-like shape of the secondary filaments of the gills, similar to those of the *Alvinella* species (Desbruyères and Laubier, 1993; Jollivet and Hourdez, 2020). The other *Paralvinella* species were subdivided in two other subgenera: *Miralvinella* and *Paralvinella* as the result of differences in their modified pairs of tentacles in males.

Recent molecular phylogenies, on the other hand, have produced conflicting results regarding the subdivision of the family into two monophyletic genera. Jollivet et al. supported this morphological view, as well as the existence of the *Nautalvinella* subgenus, based on enzyme isoforms similarities (Jollivet et al., 1995). Another study by Jollivet and Hourdez, based on a set of 278 orthologous transcripts, came to the same conclusion, with *P. p. irlandei* being a sister species to the other *Paralvinella* species (Jollivet and Hourdez, 2020). Counter to this, *Cox1* phylogenies ((Vrijenhoek, 2013), (Jollivet and Hourdez, 2020) and a recent wide phylogeny for Terebelliformia by Stiller and colleagues, based on several hundred orthologous genes derived from transcriptomic datasets, proposed with high support toward the polyphyly of the *Paralvinella* genus, with *P. p. irlandei* being sister to the other Alvinellidae (Stiller et al., 2020). However, none of them included sequences from *P. unidentata*, in contrast to studies suggesting the monophyly of the genus, which then anchor the species *P. p. irlandei*, *P. p. pandorae* and *P. unidentata* closer together in the phylogeny (Desbruyères and Laubier, 1993; Jollivet et al., 1995; Jollivet and Hourdez, 2020). It appears that the phylogenetic relationship between the three *Nautalvinella* species and the *Alvinella* genus is therefore key to understanding the deep phylogeny of the Alvinellidae.

Here, we propose a new molecular phylogeny of the family Alvinellidae based on several hundred orthologous genes derived from RNAseq datasets, notably improved by a new sequencing of the species *P. unidentata* and other species from both the western Pacific and the Indian Ocean. The phylogeny therefore includes all described species of Alvinellidae, with the exception of *P. p. pandorae*, and the rare species *P. dela* and *P. bactericola* for which no exhaustive molecular data or frozen-preserved samples exist to date. This phylogeny is rooted with species of the sister group Ampharetidae, as well as more distantly-related species of Terebellidae and the coastal pectinariid worm *Pectinaria gouldii* as outgroup, in accordance with the currently accepted phylogeny of the Terebelliformia worms (Rousset et al., 2007; Stiller et al., 2020). Finally, the results obtained by different phylogenetic methods were compared, following Maximum Likelihood approach on a gene concatenation, on the one hand, and a coalescent approach from gene trees, on the other hand.

## 2. Materials and Methods

### 2.1 Animal Collection, Sequencing and Assembly

Transcripts of *A. pompejana* were predicted from the genome assembled at the chromosomal level by R. Copley (NCBI accession number PRJEB46503, El Hilali et al. (2024)) with the Augustus webserver (Hoff and Stanke, 2013). Two individuals from 9°50N/EPR were used to obtain the genome, one given by C.G. Cary and the other one collected during the Mescal 2012 French cruise with ROV *Victor6000* and RV L’Atalante. The software Augustus was first trained on cDNA sequences provided by the MPI (Holder et al., 2013) and containing previous *A. pompejana* EST assemblies obtained from a Sanger sequencing (Genoscope project, see Gagnière et al. (2010) for details). The prediction was performed on scaffolds of more than 60 kb (95.8% of the assembled genome). The resulting coding sequences were filtered with Kraken v.2.0.9 and then Transdecoder v.5.5.0 to ensure data homogeneity among all final transcriptomes. RNAseq reads for the species *P. gouldii*, *Anobothrus* sp., *Amphicteis gunneri*, *Amphisamytha carldarei* and *Hypania invalida* were downloaded from the NCBI SRA database (accession numbers SRR2057036, SRR11434464, SRR11434467, SRR11434468, SRR5590961) (Kocot et al., 2016; Stiller et al., 2020).

An EST library was previously obtained for the species *P. sulfincola* in the framework of a Join Genome Initiative project (NCBI accession number PRJNA80027), led by P.R. Girguis and S. Hourdez (Girguis and Lee, 2006). The animals were collected on the Juan de Fuca Ridge during the jdFR oceanographic cruise in 2008, with the submersible Alvin on board of the R/V Atlantis. Reads were obtained by 454 Roche technology, and 24,702 transcripts were assembled using Newbler (Margulies et al., 2005).

Other alvinellid species were collected during several oceanic cruises from 2004 to 2019 on board of the N/O L’Atalante and using either the ROV *Victor6000* or the manned submersible Nautile with the exception of *P. palmiformis*, which was also collected with *P. sulfincola* during the same jdfR cruise on Juan de Fuca. Information about the sampling locations of the alvinellid worms collected along the East Pacific Rise are provided in Fontanillas et al. (2017). Other alvinellid species and the vent terebellid were sampled from different vent sites of the western Pacific back-arc basins during the Chubacarc 2019 cruise with the tele-manipulated arm of the ROV *Victor6000* and brought back to the surface in an insulated basket. *P. hessleri* was sampled from Fenway in the Manus basin, *P. fijiensis* from Big Papi (Manus Basin) and Tu’i Malila (Lau Basin), *P. unidentata* from Fenway (Manus Basin), *P.* sp. nov. from the volcano South Su (Manus Basin) and the vent terebellid Terebellidae gen. sp. (not yet described) at Snow Cap (Manus Basin) and Tui’Malila (Lau basin). Finally, *P. mira* was sampled from the Wocan vent field in the northwest Indian Ocean on the Carlsberg Ridge by HOV Jialong during the DY38 cruise in March 2017 (Han et al., 2021). *Melinna palmata* and *Neoamphitrite edwardsi*, which are shallow-water species, were respectively sampled in the bay of Morlaix and Roscoff, France. Total RNA extraction from flash-frozen tissue were performed with Trizol after tissue grinding using a ball mill. RNAseq libraries were produced at Genome Québ ec following a polyA purification of mRNAs, and sequenced using the Novaseq 6000 technology.

With the exception of *A. pompejana* and *P. sulfincola* species, all transcriptomes were assembled using a common procedure. The reads were first cleaned of adapters and trimmed with Fastp v.0.20 to retain reads with a phred-score above 25 for each base (Chen et al., 2018). Kraken v.2.0.9 was then used to remove reads corresponding to prokaryotic contamination (Wood and Salzberg, 2014). The reads retained at this stage were assembled *de novo* using Trinity v.2.9.1 (Grabherr et al., 2011). Finally, the assembled transcripts with an identity greater than 99% on 50 consecutive bases were assembled with cap3 v.10.2011 (Huang and Madan, 1999). Results of the sequencing, filtration steps, and assembly metrics are detailed in Table 1 of the Supplementary data, together with the associated command lines. Transcriptomes are available from the NCBI BioProject repository with the following accession numbers (*Alvinella caudata* XXX, *P. unidentata* XXX, *P. palmiformis* XXX, *P. mira* XXX, *P. hessleri* XXX, *P.* sp. nov. XXX, *P. fijiensis* XXX, Terebellidae gen. sp. XXX, *M. palmata* XXX, and *N. edwardsi* XXX). Other alvinellid RNAseq data, including that of *P. p. irlandei* are available from the Sequence Read Archive (SRA) database under accession numbers SRP076529 (Bioproject number: PRJNA325100) and SPX055399-402 (Bioproject number: PRJNA80027).

**Table 1:** Scores obtained for the 15 Alvinellidae topologies under different phylogenetic models. First and second columns: maximum likelihood on the nucleotide or amino-acid encoded supermatrices, run as one gene partition under the GTR+Γ or LG+Γ model. Third and fourth columns: quartet score of species trees established from gene trees, either from nucleotide or amino-acid encoded genes. For each method, the topology with the highest score is set to 0, and the difference

|  | Concatenation |  | Coalescence |  |
| --- | --- | --- | --- | --- |
|  | 499,036 sites<br>nucleotide | 277,900 sites<br>amino acid | 657 genes<br>nucleotide | 699 genes<br>amino acid |
| T1 | - 419 | - 582 | - 2,743 | - 16,170 |
| T2 | - 600 | - 619 | - 13,197 | - 14,886 |
| T3 | - 343 | - 410 | - 6,822 | - 6,420 |
| T4 | - 312 | - 428 | - 2,915 | - 6,627 |
| T5 | - 271 | - 480 | - 2,304 | - 16,085 |
| T6 | - 119 | - 208 | - 665 | - 9,202 |
| T7 | - 167 | - 88 | - 2,234 | - 1,019 |
| T8 | - 77 | - 93 | - 6,330 | 0 |
| T9 | 0 | 0 | 0 | - 802 |
| T10 | - 482 | - 632 | - 10,956 | - 19,554 |
| T11 | - 380 | - 521 | - 10,896 | - 13,790 |
| T12 | - 548 | - 645 | - 22,097 | - 27,472 |
| T13 | - 443 | - 408 | - 9,572 | - 16,903 |
| T14 | - 421 | - 492 | - 8,951 | - 17,059 |
| T15 | - 644 | - 703 | - 14,128 | - 21,391 |

The Open Reading Frames (ORFs) were then identified with Transdecoder v.5.5.0, firstly trained with the 500 longest sequences accessible from each transcriptome. The predicted ORFs were retained, stripped of the 3’ and 5’ UTR regions. The metrics associated with these different steps are also presented in Table 1 of the Supplementary data.

In total, 25 transcriptomes were assembled and compared, from 19 different Terebelliformia species including 11 species of the family Alvinellidae.

### 2.2 Search for orthologous genes and Bioinformatic Processing

Orthologous sequences were determined from the assembled transcriptomes using Orthograph v.0.7.2 (Petersen et al., 2017). Following the recommendations of Orthograph, a reference database of 1997 orthogroups (OG) was first constructed from the ODB9 database (Zdobnov et al., 2021). These OGs correspond to single copy genes in all lophotrochozoan species available when downloading ODB9 (*Lottia gigantea*, *Crassostrea gigas*, *Biomphalaria glabrata*, *Capitella teleta*, *Helobdella robusta*). Then, each Terebelliformia transcriptome was blasted against these OGs with Orthograph to identify orthologous genes. These steps attempted to identify all single-copy orthologous genes present in Terebelliformia. The results of this identification process are detailed in Supplementary data Table 1. Finally, to improve the phylogenetic signal between alvinellid worms, we retained only OGs that contained at least one sequence for the 4 most divergent clades of alvinellid species: *P. pandorae irlandei*, *P. unidentata*, *Alvinella* spp and *Paralvinella* spp. The presence of these sequences is considered mandatory.

The nucleotide sequences were aligned for each gene with MACSE v.2.05, which is a codon-gaps aligner for coding sequences (Ranwez et al., 2011), and the translated amino acid sequences were subsequently aligned with Probcons v.1.12 (Do et al., 2005). From these sequence alignments, we obtained one set of 657 OGs, in nucleotides, and one set of 699 orthologous amino-acid translated genes, present in at least 20 transcriptomes. For nucleotidic sequences, aligned fragments shorter than 60 nucleotides between two subsequent gaps were considered unreliable and discarded. The third nucleotide of each codon was also removed to reduce the saturation of the phylogenetic signal. Sequences shorter than 20% of the length of the total gene alignment were also discarded. Finally, each OG alignment was tested with IQ-TREE 2.0.3 (Minh et al., 2020) to guarantee that the phylogenetic assumptions about residue homogeneity and stationarity were not violated using the tests of symmetry at a 5% threshold (Naser-Khdour et al., 2019). When an orthologous group failed the test, biased sequences (excluding mandatory species) were eliminated until the alignment did not reject the homogeneity and stationarity hypothesis; For amino-acid sequences, filtering conditions were those described for the nucleotidic sequences, but with the removal of fragments shorter than 20 amino-acids when included between two subsequent gaps. The slight difference in the filtering of sequences for residue homogeneity and stationarity, encoded as nucleotides or amino acids, led to the difference in the number of OGs finally selected for the two sets of sequences in the phylogenetic reconstruction.

These sets were then concatenated into two supermatrices. The first, in nucleotides (two first codon positions of the coding sequence), contained 657 genes and 499,036 sites, with *A. caudata* being the longest sequence with 461,042 sites (92% completeness), and *P. gouldii* the shortest with 245,846 sites (49% completeness). The second, in amino-acids, contained 699 genes and 277,900 sites, with *A. caudata* being the longest sequence (255,557 sites, 92% completeness) and *P. gouldii* the shortest (139,067 sites, 50% completeness). The length and completeness of each species concatenated sequence is given in supplementary Table 2.

### 2.3 Phylogenetic inference

#### 2.3.1 Tree reconstructions with the identification of well-resolved clades

A global unrooted phylogeny was evaluated from the supermatrices with the maximum likelihood (ML) approach implemented in IQ-TREE 2.0.3. The supermatrices were first partitioned according to each gene, and we used the merging strategy implemented in IQ-TREE to retrieve relevant merged partitions (Chernomor et al., 2016). For each merged partition, ModelFinder, was used to infer the best-fitting evolutionary model according to the minimized Bayesian Information Criteria (BIC) (Kalyaanamoorthy et al., 2017). For nucleotides and amino-acids sequences, heterogeneity of the evolutionary rate between sites was modelled by a Γ distribution with four discrete classes (Yang, 1994). For amino-acid sequences, empirical or ML equilibrium amino-acid frequencies were both tested, and mixture models were also added to the standard test of ModelFinder (Si Quang et al., 2008; Le et al., 2008, 2012). The reconstructed trees were then secondarily rooted using the pectinariid sequences of *P. gouldii* as the outgroup using the software FigTree v.1.4.4. Finally, the robustness of the tree was inferred *via* 1,000 resamplings of the datasets with bootstraps. The unrooted phylogeny was also obtained in the coalescent framework. To this end, gene trees, for either nucleotide or amino-acid encoded sequence alignments, were first optimized using IQ-TREE. Substitution models were tested with ModelFinder, including mixture models for amino-acid sequences. Species trees were obtained from gene trees using the supertree method implemented by ASTRAL v.5.7.8 (Zhang et al., 2018), which is assumed to be consistent under the multispecies coalescent model. Branches with low bootstrap support in the gene trees (*<*10%) were contracted beforehand, according to the recommendations of the authors. The robustness of the tree was inferred *via* 1,000 resamlings of the gene trees datasets with bootstraps.

#### 2.3.2 Tree reconstruction with a gene-to-gene approach and the coalescent method Testing alternative topologies with the AU test

It appeared that the two most ancestral nodes at the root of the Alvinellidae family were the most conflicting ones depending on the reconstruction methods or the genes used in the reconstruction. However, both the *Paralvinella* clade (excluding *P. p. irlandei* and *P. unidentata*) and the set of outgroup species were well resolved in the first step of the tree reconstructions, whatever the method used. Reconstructions therefore led to the co-occurrence of 4 distinct alvinellid clades that can be arranged in 15 alternative topologies, some which being more probable than the others.

In order to test whether the most likely topology really was the most robust one, we evaluated the confidence of these 15 topologies using the AU test of IQ-TREE in a ML framework, namely the GTR+Γ4+F model for the nucleotide-encoded supermatrix taken as a single partition, or the LG+Γ4+F model for the amino-acid encoded supermatrix also taken as one partition. The evolutionary models used for the two supermatrices correspond to the most commonly chosen models obtained under a complete gene partition. The AU test was used at a 5% threshold with 10,000 RELL replicates to compare alternative topologies against the most likely topology obtained for each supermatrix (Shimodaira, 2002). This test gives the probability, based on p-values, that an alternative topology has a significantly better likelihood than the reference topology, after a site-to-site resampling of the supermatrix.

We also transposed the AU test in the coalescent framework to test for the significance, based on p-values, that an alternative species tree topology has a higher quartet score than the reference species tree topology after gene trees resampling. The multiscale bootstrap sampling procedure described in Shimodaira (2002) was thus recoded using genes with 10 successive resamplings of the gene datasets with 10,000 bootstraps, each comprising from 0.5 to 1.4 times the total number of genes to take into account for the effect of the number of genes used in the reconstruction. The ML parameters of the method were estimated as described in Shimodaira (2002), and scripts for this procedure are given in Supplementary material. Again, topologies were compared to the best-scoring species tree topology obtained with nucleotide or amino-acid-encoded gene-trees at a 5% threshold.

### Accounting for ILS and GTEE using the 15 constrained tree topologies

Gene tree discordance was assessed in an attempt to distinguish between gene tree estimation error (GTEE), incomplete lineage sorting (ILS) or gene flow between species. For each gene, the 15 candidate topologies were evaluated under ML using the best evolutionary model according to ModelFinder. Topology *k* was then given a Bayesian posterior probability from its likelihood *L*_k_ using the approximation.

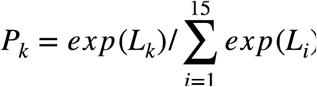

In the case low ILS, one would expect to sample randomly between the 15 candidate topologies for genes with a low number of phylogenetically informative sites due to GTEE, while only one or a few preferred topologies should prevail for genes with a high number of informative sites. On the contrary, high ILS would imply that gene tree randomness remains high, independently of the load of the phylogenetic signal. Therefore, genes were split into four categories according to their number of parsimony-informative sites for the nodes at the root of the family Alvinellidae in order to observe whether gene tree discordance was similar across the different categories. The deviation of the gene tree distribution from randomness is tested by the *χ*2 statistics against a multinomial equiprobable probability.

To test whether allele introgression between the oldest ancestors also occurred, the gene tree topologies’ probabilities were mapped on a ternary plot, allowing us to evaluate the number of genes that are strongly associated with one particular topology (*Pr >* 0.8). In that case, the 15 topologies were collapsed into three possibilities corresponding to different quartets at the root of the Alvinellidae family: ((*P. p. irlandei,P. unidentata*), (*Alvinella*, *Paralvinella*)), or ((*P. p. irlandei*, *Alvinella*), (*P. unidentata*, *Paralvinella*)), or ((*Alvinella*, *P. unidentata*), (*P. p. irlandei*, *Paralvinella*)). In case of a strong allele introgression between ancestral lineages in the early steps of the Alvinellidae radiation, one would expect to see one quartet supported by a majority of genes (corresponding to the species tree), and a second quartet supported by a lower but substantial number of genes (as a consequence of the between-species gene flow) (Cai et al., 2021). If no introgression occurred, the quartet corresponding to the species tree should be the most supported one, while the alternative quartets should have the same prevalence among gene trees. The imbalance of the distribution of gene trees between two quartets is tested against a binomial law.

Finally, we expect gene tree estimation using the 15 constrained topologies to reduce the overall GTEE compared to its free-running counterpart. As coalescent methods are very sensitive to GTEE (Roch and Warnow, 2015; Nute et al., 2018), we estimated the species tree with ASTRAL based on the constrained gene tree estimation, with gene tree topologies being drawn from their Bayesian posterior probability to take into account their uncertainty. This procedure was conducted on both nucleotide-encoded genes and amino-acid encoded genes. To summarize the information coming from the two types of sequences, a global species tree score was calculated. Nucleotide normalized quartet scores *N*, comprised between 0 and 1, were combined with amino-acid normalized quartet scores *A*, between 0 and 1, by considering the Euclidean distance *d* between pairs of sequence datasets (*N, A*) and a hypothetical ideal topology scoring (1, 1):

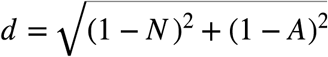

This topology would correspond to an ideal tree that would maximize the quartet scores both for nucleotide and amino-acid encoded genes. Thus, the lower the distance estimated, the better the topology.

The distance obtained for each candidate topology was then tested against the best-scoring topology with our transposed AU test at a 5% p-value threshold with resamplings of 10,000 gene trees drawn according to their Bayesian posterior probability.

#### 2.3.3 Molecular dating of the family’s radiation

The alvinellid phylogeny was dated with the software Phylobayes v.4.1. (Lartillot and Philippe, 2004; Lartillot et al., 2007). The uncorrelated Gamma model (Drummond et al., 2006), Cox-Ingersoll-Ross(CIR) model (Lepage et al., 2007) and the log normal model (Thorne et al., 1998) were tested on the concatenated amino acid-encoded genes for the most relevant species tree topology. Considering than Terebelliformia fossils are the most abundant in the Carboniferous/ Devonian time (Sepkoski Jr., 2002), the age of the tree’s root (divergence between *P. gouldii* and other terebellid, ampharetid and alvinellid species) is given by a Gamma distribution *prior* with a mean of 330 million years and a standard deviation of 200 million years. We also used two calibration points. The first one corresponded to the divergence between the species *P. palmiformis* and *P. grasslei*, estimated to be 23 to 34 millions of years ago (Ma) as a consequence of the subduction of the Farallon plate under the North American plate (Tunnicliffe, 1988)), and the second one to the divergence between the two geographic forms of the species *P. fijiensis* sampled in the Manus and Lau back-arc basins, which opening is estimated between 2 and 4 Ma (Desbruyères et al., 2006; Boulart et al., 2022). Finally, we tested the hypothesis of a more ancient *prior* on the age of the tree’s root (420 millions years, standard deviation 100 millions years) according to the oldest fossils attributed to terebellids (Thomas and Smith, 1998; Little et al., 1999; Vinn and Toom, 2014).

## 3. Results

### 3.1. Global phylogenetic inference

A free-running phylogeny was performed under a partition gene model on both the nucleotidic and amino-acid supermatrices, or using the coalescent method on the single gene datasets, based on either nucleotide or amino-acid encoded sequence alignments. Both approaches identify three main clades inside terebellomorph species, which are the Alvinellidae family, the Ampharetidae family comprising *Anobothrus* sp., *H. invalida*, *A. carldarei* and *A. gunneri*, and a third group composed of the Terebellidae species (Terebellidae gen. sp. and *N. edwardsi*) and the melinnid species (*M. palmata*), as shown in Figure 1. *M. palmata* is a sister species of terebellidae species for all methods with the exception of the phylogeny obtained from the nucleotide-encoded supermatrix, where it represents a sister group to the three polychaete families (Alvinellidae, Terebellidae and Ampharetidae excluding the outgroup *P. gouldii*). *Anobothrus* sp. also represents a sister species to Ampharetidae species in phylogenies using the amino-acid encoded sequences (bootstrap value of 92% for the supermatrix approach but only 51% in the coalescent method), but is sister to other Ampharetidae and Alvinellidae species using nucleotide-encoded sequences with a greater robustness (with a bootstrap value of 99% and 95%, respectively). All phylogenies with their node supports are included in the supplementary data, Figure 1 to 4. In further analyses, we chose to only consider phylogenies that group *M. palmata* in the Terebellidae clade and separate Alvinellidae and Ampharetidae as two monophyletic families (Fig. 1), according to current views on the Terebelliformia phylogeny (Stiller et al., 2020).

**Fig. 1:**
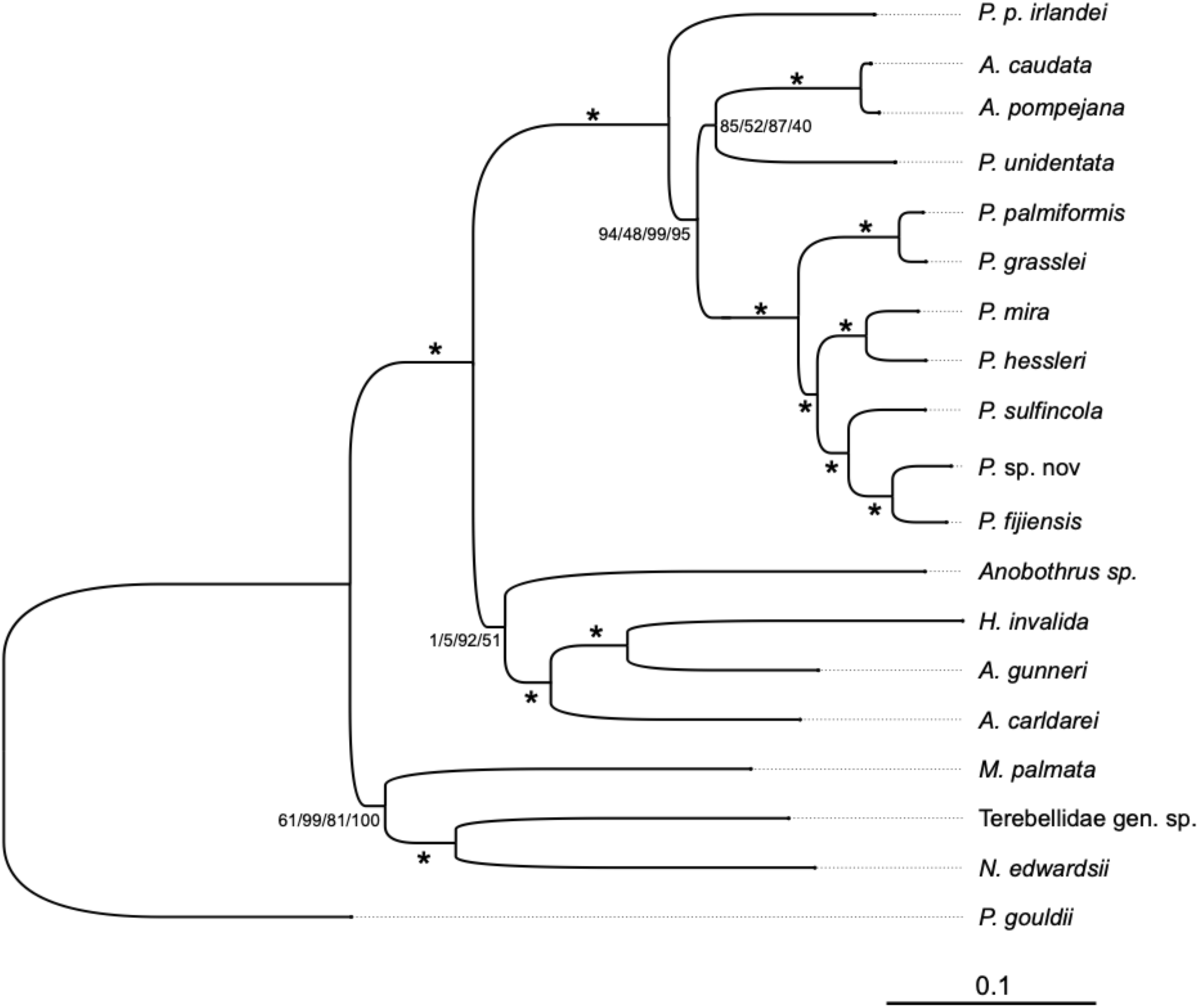
Best-scoring phylogeny according to different phylogenetic inference methods, with *P. p. irlandei* sister to the other alvinellid species and clustering of *P. unidentata* with the *Alvinella* species. Branch lengths are optimized under a partitioned model from concatenated amino-acid encoded supermatrix. Node supports are given in the following order: site sampling from nucleotide-encoded supermatrix, gene sampling with coalescence model based on nucleotide encoded genes, site sampling from amino acid-encoded supermatrix, gene sam-pling with coalescence model based on amino acid encoded genes. (*) indicates bootstraps of 100% for all phylogenetic methods.

Within the Alvinellidae, all methods give a maximum bootstrap value to the grouping of the two *Alvinella* species. In the *Paralvinella* genus, with the exception of *P. unidentata* and *P. p. irlandei*, the phylogeny of species is also resolved without ambiguity, with a maximum of confidence on all nodes regardless of the phylogenetic method used.

Conversely, the placement of the species *P. unidentata*, and more importantly *P. p. irlandei*, is not well supported. Depending on the number of genes considered (with and without taking into account the mutational amino-acid bias observed between sequences), the evolutionary model used, or the types of sequences, the placement of these two species in relation to the well-defined groups of the other *Paralvinella* species on one hand, and the two species of the genus *Alvinella* on the other hand, was fluctuating at the root of the family Alvinellidae. By bootstrapping sites on the concatenated set of genes in the ML approach or the genes themselves in the coalescent approach, the best supported topology was however the topology placing *P. p. irlandei* as a sister species to all alvinellid species, including *Alvinella* spp. as shown in (Figure 1, 1) on the consensus trees obtained by the different methods. However, it is worth-noting that only 10.5% of nucleotide-encoded genes and 11.0% of amino-acid encoded genes conformed this topology, named T9 in figure 2. By comparison, 10.0% and 10.3% of genes favoured the monophyly of the *Paralvinella* genus with the grouping of *P. p. irlandei* and *P. unidentata*, named T6 in figure 2. Because of this uncertainty, and to better explore the evolutionary processes that could have occurred during the radiation of the Alvinellidae, we chose to explore more specifically the confidence in the resolution of these deep nodes. To this end, we fixed the topology obtained for both the outgroups and the other *Paralvinella* species, as they were well resolved, in order to perform AU tests which are best suited to provide confidence in the robustness of the selected tree.

**Fig. 2:**
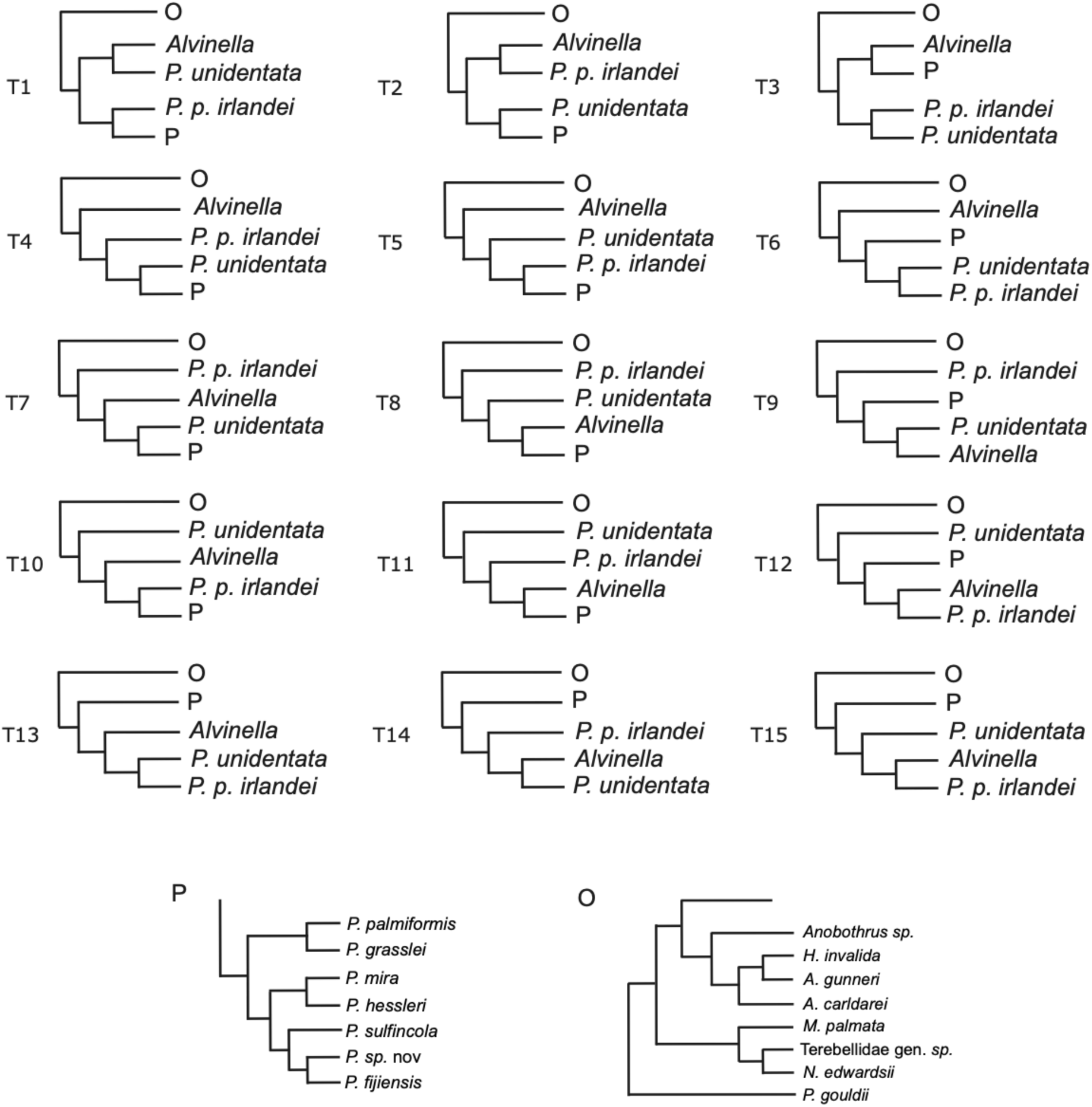
Constrained topologies. The topologies only differ in the arrangement of the species *P. unidentata*, *P. p. irlandei*, when compared with the genera *Paralvinella* and *Alvinella* at the root of Alvinellidae. The *Alvinella* group comprises the species *A. pompejana* and *A. caudata*, and the *Paralvinella* group (P) comprises all species *Paralvinella* except *P. p. irlandei* and *P. unidentata*. Trees are rooted with a set of outgroups (O) with the same species arrangement comprising species from other closely-related families of Terebelliformia (Terebellidae+Ampharetidae+*P. gouldii*).

### 3.2 Evaluating bifurcations at the root of Alvinellidae

In order to assert the alvinellid phylogeny at the root of the family, we exhaustively compared the likelihoods obtained for the 15 possible tree topologies when both the outgroup subtree (Terebellidae+Ampharetidae rooted with *P. gouldii*) and the *Paralvinella* subtree (excluding *P. p. irlandei* and *P. unidentata*), considered as resolved, are constrained. Figure 2 lists all these topologies, numbered from 1 to 15.

The results of the AU tests for the 15 tree topologies using our two sequence datasets (CDS or translated proteins) are presented in Table 1. From these analyses, four topologies stand out:

- Topologies 7, 8 and 9, in which *P. p. irlandei* is sister to the all other Alvinellidae. In topology 7, the *Alvinella* lineage is closer to *P. p. irlandei*, topology 8 brings *P. unidentata* closer to other *P. p. irlandei*, and in topology 9, *A. pompejana*, *A. caudata* and *P. unidentata* are sister species. According to the AU test, the best-ranking topology T9 obtained in the different phylogenetic methos was not significantly better than the other topologies T8 and T7, except T7 which was significantly worse than T9 in the nucleotide+ML analysis (p-value=0.3%). Following this testing, we finally retained T9 as the most appropriate topology over the three;
- Topology 6, in which the *Alvinella* lineage is sister to all *Paralvinella*, including *P. p. irlandei* and *P. unidentata*. According to the AU test, the best-ranking topology T9 (using the ML inference on nucleotide sequences), or the topologies T9 and T8 (using the coalescence approach with both nucleotide and amino acid-encoded genes) were not significantly better than T6. However, T6 was significantly worse than T9 in the amino acid+ML approach (p-value=2.1%).

### 3.3 Phylogenetic discordance between genes

We then focused on the evaluation of the phylogenetic signal incongruousness at the gene level for the root of the family. We estimated the likelihood of the 15 constrained topologies for each gene separately, either on their coding sequences or their translated protein counterparts (and depending on their number of parcimony-informative sites, see supplementary figure 5). As shown in Figure 3b. and 3d., all topologies were supported by a high proportion of genes regardless of the strength of the phylogenetic signal. The distribution only differed from equiprobable between the 15 topologies for highly-informative nucleotidic encoded genes (*χ*2, *p*−*value* = 3.1%). Interestingly, genes containing more parsimony-informative sites, and thus having more power to distinguish between topologies, did not exclude any topology but the topologies T6 and T9 were supported by a higher proportion of genes than 1*/*15 for nucleotide-encoded genes (*p*−*value* = 0.6% and 0.3% respectively) and amino-acid encoded ones (*p*−*value* = 1.5% and 0.7%). As a consequence, despite T6 and T9 appeared more often in the gene trees, the discordance between gene histories was attributable to ILS, and not solely to random gene tree estimation error and a loss of phylogenetic signal at some slow-evolving genes in which a higher GTEE is prone.

**Fig. 3:**
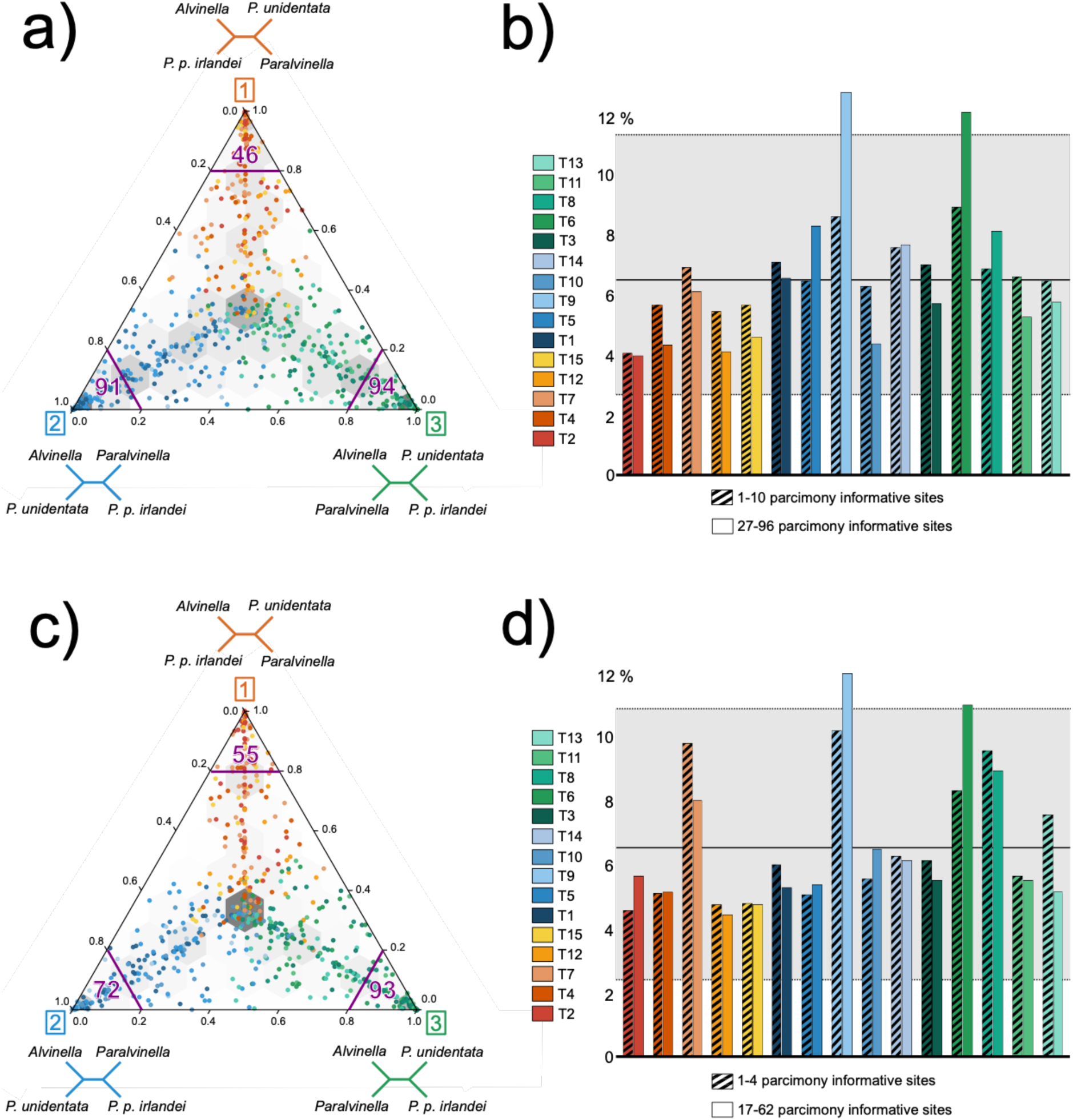
Topology weight across genes. (a) Nucleotide-encoded gene associations with the three possible scenarios labelled 1, 2 and 3. Number of genes strongly associated with one scenario (*Pr >* 0.8) are indicated in each pole. (b) Distribution of genes’ proportions according to the number of parsimony-informative sites for nucleotide sequences. Greyed area represents the 96% interval for random binomial distribution (1*/*15 probability of choosing one topology). (c) Amino acid-encoded gene associations with the three possible scenarios labelled 1, 2 and 3 at the root of the family. Number of genes strongly associated with one scenario (*Pr >* 0.8) are indicated in each pole. (b) Distribution of genes’ proportions according to the number of parsimony-informative sites for amino-acid sequences. Greyed area represents the 96% interval for random binomial distribution (1*/*15 probability of choosing one topology).

In Figure 3a. and 3c., we collapsed the 15 constrained topologies to the three possible scenarios labelled 1, 2 and 3, to identify a potential bias of support in the shared gene histories between the *Alvinella*, *Paralvinella*, *P. unidentata* and *P. p. irlandei* lineages. Indeed, one of the three scenarios should account for the true species tree, and two alternative scenarios should be the result of either introgression, ILS or GTEE (Cai et al., 2021). Moreover, observing a frequency bias in the sampling of the two alternative scenarios is the result of specific introgression between two ancestors. We tested this bias by considering the null hypothesis that the two alternative quartets were equally sampled in gene trees, following a binomial distribution of parameters *n*, being the number of genes falling in the alternative quartets, and *p* = 0.5, the probability of choosing one quartet over the other (Pease et al., 2018). Considering nucleotide-encoded genes, 182 genes favor scenario 1 with a closer relationship between *Alvinella*+*P. p. irlandei* opposed to *P. unidentata* +*Paralvinella* (55 genes assigned with *Pr >* 0.8), 234 genes favor scenario 2 in which *Alvinella*+*P. unidentata* is opposed to *Paralvinella*+*P. p. irlandei* (91 with *Pr >* 0.8) and 241 genes are supporting scenario 3 where *Alvinella*+*Paralvinella* is opposed to *P. p. irlandei* +*P. unidentata* (94 with *Pr >* 0.8). For amino-acid encoded genes, these scenarios are respectively supported by 205 (55), 234 (72) and 260 (94) genes. For nucleotide-encoded genes, choosing scenario 2 or 3 as the true species tree results in scenario 1 being less supported than the alternative topology (scenario 1 against 2: *X* ∼ *B*(416, 0.5), *Pr*(*X* ⩽ 182) = 0.006, scenario 1 against 3: *X* ∼ *B*(423, 0.5), *Pr*(*X* ⩽ 182) = 0.002).

On the contrary, scenario 2 and 3 are not distinguishable if scenario 1 in the true topology (*X* ∼ *B*(475, 0.5), *Pr*(*X* ⩽ 234) = 0.39). For amino-acid encoded genes, choosing scenario 2 or 3 as the true species tree results in scenario 1 being less supported than the alternative topology 3 (*X* ∼ *B*(465, 0.5), *Pr*(*X* ⩽ 205) = 0.006) or marginally less supported than scenario 2 (*X* ∼ *B*(439, 0.5), *Pr*(*X* ⩽ 205) = 0.09). Again scenario 2 and 3 are not distinguishable if scenario 1 in the true topology (*X* ∼ *B*(494, 0.5), *Pr*(*X* ⩽ 234) = 0.13). Consequently, one of the scenarios 2 or 3 is likely the true species tree while the second is the result of massive introgression between ancestors of the *P. pandorae* and *P. unidentata*. In this case, gene trees supporting scenario 3, which brings closer the *Alvinella* species and *P. p. irlandei*, are the result of ILS and potential GTEE.

### 3.4 Testing for the right species tree topology with constrained gene trees and the coalescent approach

In our final approach, we attempted to reconstruct the species tree using the coalescent method from gene trees constrained to one of the 15 constrained topologies. The goal was to reduce the overall gene tree estimation error (GTEE), assuming that incomplete lineage sorting (ILS) and inter-species gene flow were low at all nodes of the tree with the exception of the two nodes at the root of the family Alvinellidae. When evaluating the 15 topologies with this approach (see: Figure 4a.), T7 had on average the highest quartet scores both for nucleotide-encoded genes and their translated amino-acid counterparts, although this result was sensitive to the resampling of genes and gene trees uncertainty. Figure 4b. displays the combined quartet scores obtained by each topology as the Euclidean distance to a hypothetical ideal topology at coordinates (1,1) that would maximize the quartet scores for both types of sequences. In this representation, T7 also obtained the best combined score, although not statistically better than T8 and T9 (*p−value* = 0.23, 0.34 respectively). In all these topologies, *P. p. irlandei* is paraphyletic with the other *Paralvinella* species. The relationship of *P. unidentata* with other alvinellid clades is however different in these three topologies, being sister species to *Paralvinella* in T7, to *Alvinella* in T9 or to *Alvinella*+*Paralvinella* in T8. The fact that T7 has the greatest scores may however be explained by its intermediate topology between T6 and T9, which were two most frequent co-occurring topologies in the gene trees sets (see Discussion, Paraphyly of the genus *Paralvinella*).

**Fig. 4:**
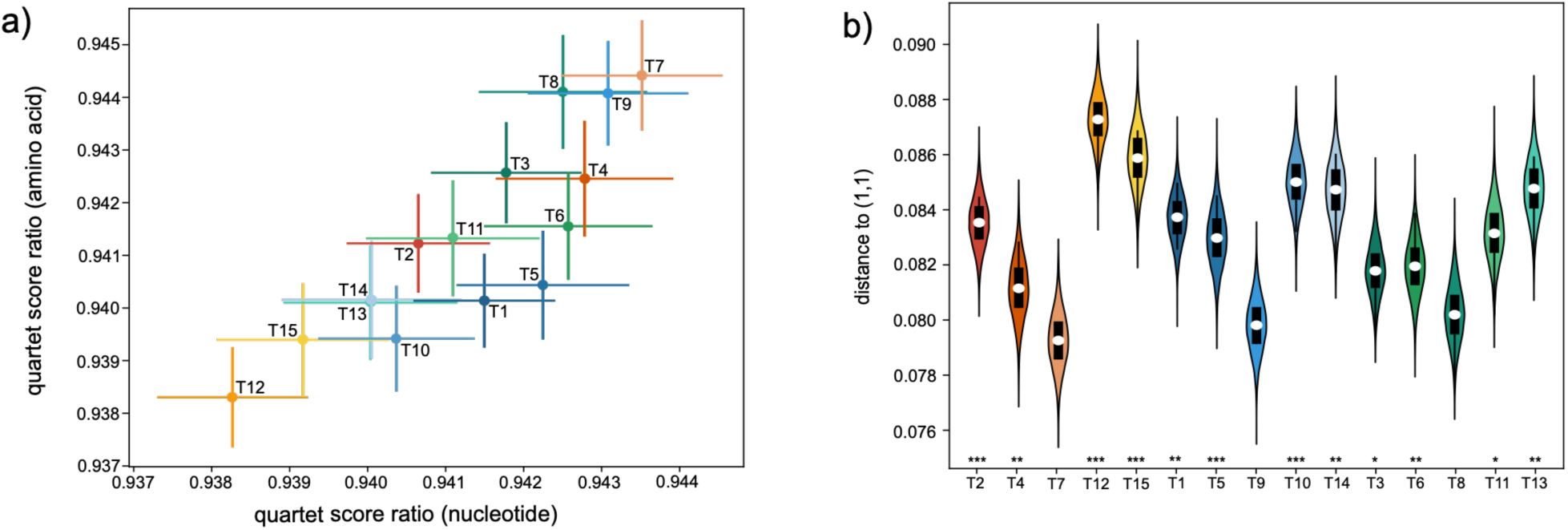
Fitting scores of the species tree topology by ASTRAL from the 15 constrained topologies, with standard deviations. Each evaluation is the result of 10,000 resamplings of gene trees. (a) biplots of quartet scores (in %) obtained for each topology from the amino acid and nucleotide sequence datasets. A score closer to 1 indicates that the species-tree topology is more in agreement with the set of gene trees. (b) Euclidean distance *d* estimated for each topology to the point (1,1), which represents an ideal case where all gene trees agree with the proposed species tree topology. Each topology TX is tested against the best scoring topology T7. The hypothesis *d*_7_ *< d_X_* is tested with a transposed AU test based on gene re-sampling. If the test fails, then T7 is not better than TX: *\** : *p−value <* 5%, *\*\** : *p−value <* 1%, *\*\*\** : *p−value <* 0.1%.

### 3.5 Molecular dating of the alvinellid radiation

We estimated the age of the alvinellid worm’s radiation under the T9 topology hypothesis. Using 330 Ma as a prior for the emergence of Terebelliformia, the three models used for molecular dating (Log-normal, CIR, and uncorrected Gamma multipliers) gave different results about the age of the radiation. The log-normal model proposed 51 millions of years ago (Ma, 95% confidence interval: 42-65), however it appeared to significantly underestimate ages since the root of the tree was estimated to be 85 millions of years old (My, confidence interval: 70-109). This is far from our expectations based on the age of abundant terebellomorph fossils from the Devonian-Carboniferous period (Sepkoski Jr., 2002). On the other hand, the uncorrelated Gamma multipliers model, which assumes that the evolutionary rates of branches are uncorrelated with time, placed the age of the tree’s root (Terebelliformia radiation) at 292 My (142-504 Ma), which is very close to the expected age. The radiation of the Alvinellidae was then estimated at 99 Ma (56-180 Ma). However the 95% confidence intervals remained very large with this model. Finally, estimates proposed by the CIR model had narrower confidence intervals and are shown in Figure 5. The age of the terebelliform MCRA is 160 My (135-198 My), which is still younger than the age of the oldest fossils attributed to terebellomorphs. With this method, the age of the Alvinellidae radiation was however very close to that of the uncorrected Gamma multipliers model with an estimated age of 91 Ma (77-112 Ma). Increasing the *prior* for the age to the emergence of terebellid-like worms to a mean of 420 Ma did not affect much the age of the alvinellid radiation with an estimated date of 86 Ma (72-105 Ma) for the CIR model and 113 Ma (70-181 Ma) for the uncorrelated Gamma multipliers model. In the two last models, the radiation of the Alvinellidae was a rapid event occurring in the Late Cretaceous. The estimates obtained with the CIR model for alternative topologies T6 and T7 are given in supplementary data, Figure 6.

**Fig. 5:**
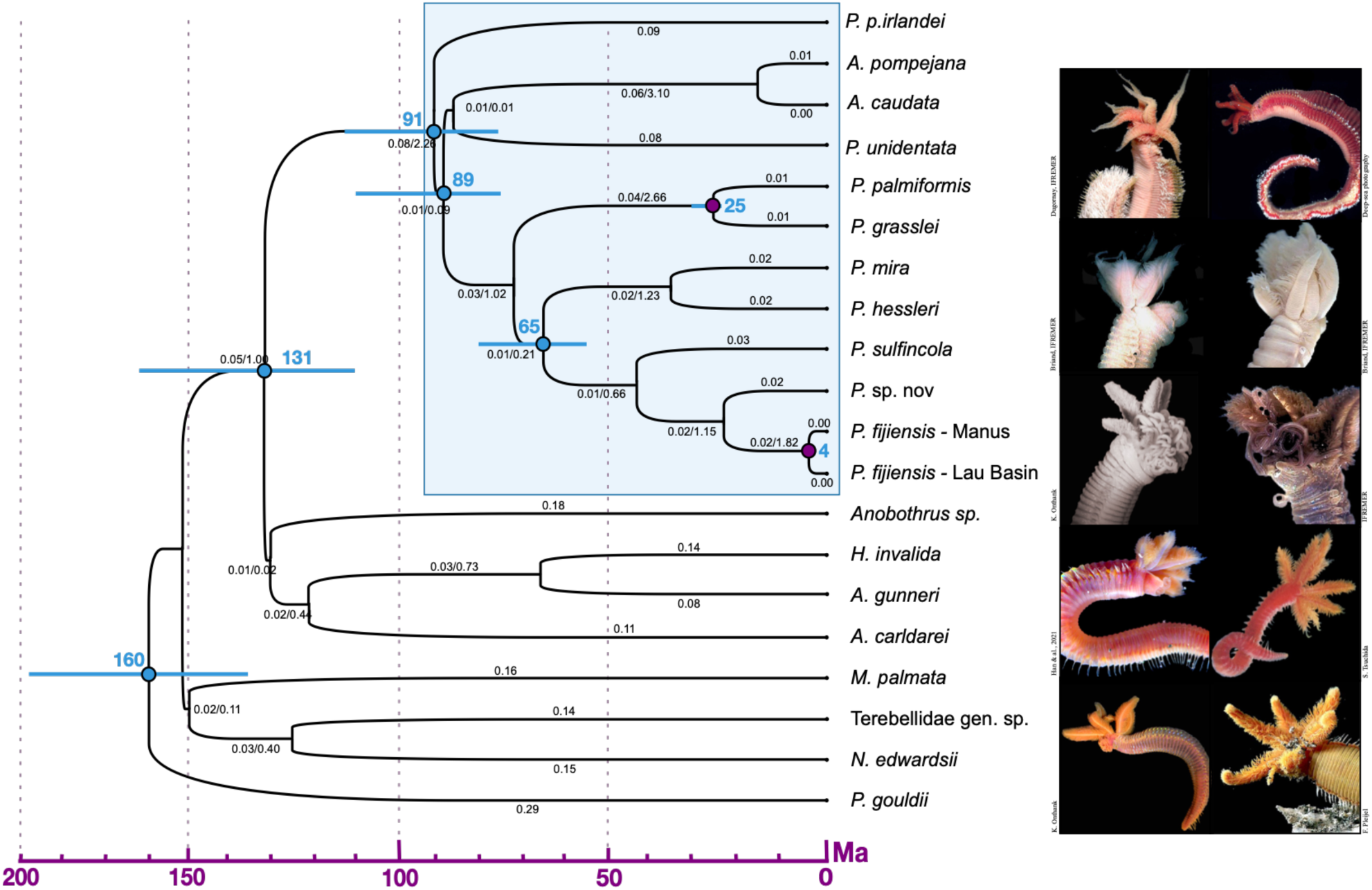
Chronogram of the family Alvinellidae under the CIR model with two calibration points (in purple), assuming T9 topology. Branch lengths are reported in expected numbers of amino-acid substitutions (LG+Γ) and coalescent time unit (obtained from unconstrained gene trees). The ages of some nodes of interest with 95% confidence intervals are reported in millions of years ago (Ma). Illustrations: (A) *A. pompejana*, (B) *A. caudata*, (C) *P. uniden-tata*, (D) *P. p. irlandei*, (E) *P. palmiformis*, (F) *P. grasslei*, (G) *P. mira*, (H) *P. hessleri*, (I) *P. sulfincola*, (J) *P. fijiensis*.

## 4. Discussion

### 4.1 Paraphyly of the genus Paralvinella

The initial description of the family by Desbruyères and Laubier subdivided the family into two genera, mainly based on the shape of the gills, the position and shape of the ventral uncini and the position of the first setiger (Desbruyères and Laubier, 1986; Jouin and Gaill, 1990; Jollivet and Hourdez, 2020). These authors also proposed a taxonomic arrangement in which *P. unidentata* was closer to the sibling pair of species *P. p. irlandei* and *P. p. pandorae* within the *Paralvinella* genus, and subsequently grouped into the *Nautalvinella* subgenus (Desbruyères and Laubier, 1993). This grouping proposal was equivalent to the T6 topology, which is not the best topology identified by the molecular phylogeny of Alvinellidae proposed here, but is also well supported by individual gene trees when constrained among the 15 most relevant tree topologies, even for the most phylogenetically-informative genes. By contrast, topologies T7, T8 and T9, within which *P. p. irlandei* is a sister species to the other Alvinellidae, got the best overall scores (maximum likelihood or super-tree resolution). These topologies, notably T9, are also well supported by a high number of genes, suggesting that the genus *Paralvinella* is paraphyletic. This contrasts with the initial view of a family split into two genera with *Alvinella* species sister to *Paralvinella* species. The paraphyly hypothesis of *Paralvinella* agrees with previous published phylogenies based on either mitochondrial *Cox1* sequences or sets of orthologous sequences derived from transcriptomes, combined with some morphological characters (Vrijenhoek, 2013; Stiller et al., 2020; Jollivet and Hourdez, 2020). Following this result, we can now state that the two *P. pandorae* sub-species must belong to a new alvinellid genus, sister to other alvinellid species, which can be named as *Nautalvinella*. The positioning of this genus may fit well with the opportunistic way of life of these species which also display less adapted gills (smaller volumes of gaz exchanges) and a morphology closer to ampharetid worms (D. Jollivet, pers. obs.). The positioning of *P. unidentata* is however much more uncertain. Topologies T7, T8 and T9 indeed differ in the positioning of this species in relation to the rest of the Alvinellidae. In T7, *P. unidentata* is closer to other *Paralvinella* species, while it is closer to *P. p. irlandei* in T8, and closer to *Alvinella* in T9. It is worth noting that the grouping of *P. unidentata* and *Alvinella* (T9) is in agreement with some traits concerning the gills’ morphology: in particular, the filaments of *P. unidentata* are flattened, which is unique compared to other *Paralvinella* species, and similar to the lamellar shape of the gills of the two *Alvinella* species. This topology thus raises the question of the potential inclusion of *P. unidentata* inside the *Alvinella* genus. However, the overall structure of the gills of *P. unidentata*, with a tip devoid of filaments and comb-like inserted secondary filaments, brings the species closer to *P. p. irlandei* (Desbruyères and Laubier, 1993). The high variation of the shape and size of Alvinellids’ gills is likely due to natural selection, as they are crucial to enduring the long periods of hypoxic conditions associated with the vent habitat, while being able to rapidly capture massive amounts of oxygen in well-oxygenated water when the worm is no longer exposed to the fluid mixing (Jouin and Gaill, 1990). As such, the shape of the gills can represent a valid morphological criterion to separate species between the three alvinellid genera if the *Nautalvinella* is elevated to the genus level.

Considering that the paraphyly of *Paralvinella* is the preferred solution, we chose to retain T9, as shown in Figure 5, as the most reliable species tree topology for producing a chronogram. The radiation of the Alvinellidae is undoubtedly a difficult case to resolve where multiple short branches (divergence *<* 1%) are buried deep in the tree topology. The radiation leading to the separation of the *Alvinella*, *P. unidentata*, *P. p. irlandei*, and other *Paralvinella* lineages is likely a fast event in the timescale of the whole family, and possibly linked to some adaptive strategies to cope with different thermal habitats. This also agrees well with the idea that the evolutionary rate may be slow in alvinellid worms (Chevaldonn é et al., 2002; Fontanillas et al., 2017), consequently maintaining ancestral polymorphisms for a very long time (Fontanillas et al., 2017). For example, in *P. unidentata* and *P. fijiensis*, present-day populations that have been separated for 3 to 4 million years also display short coalescent times, between 0.05 and 0.35 (see supplementary data Fig. 1), and Jang et al. (2016) estimated the split of the the northern East Pacific Rise and the northeastern Pacific Antarctic Ridge populations to about 4.2 million years ago for *A. pompejana*, despite the maintenance of shared polymorphisms between the two metapopulations.

### 4.2 Revision of the genus Paralvinella

Desbruyères and Laubier proposed to divide the genus *Paralvinella* into three subgenera on the basis of morphological characters (Desbruyères and Laubier, 1993):

- *P. Paralvinella*, which groups the species *P. palmiformis*, *P. grasslei*, *P. fijiensis* and *P. sulfincola*;
- *P. Miralvinella*, which groups the species *P. hessleri*, *P. dela* and *P. bactericola*;
- *P. Nautalvinella*, which includes the species *P. unidentata*, *P. p. irlandei* and *P. p. pandorae*.

Several criteria were used to differentiate these subgenera, notably the presence of a pair of specialized sexual tentacles (used in the male-female pairing during reproduction) on the buccal apparatus of males in *P. Paralvinella* (three-lobed) and *P. Miralvinella* (pointed and coiled) and absent in *P. Nautalvinella*, the shape of the gills (filaments on the opposite sides in *P. Paralvinella* and *P. Miralvinella*, and comb-like in *P. Nautalvinella*), as well as the presence of digitiform notopodial lobes on the anterior part of *P. Paralvinella* and *P. Miralvinella* which are absent in *P. Nautalvinella*. Other morphological criteria were used to differentiate these subgenera despite being more variable, such as the number of segments of the animals (55 to 180 setigers in *P. Paralvinella versus* 50 to 75 in *P. Miralvinella* and *P. Nautalvinella*), or the position of the first uncini (setigers 12 to 26 in *P. Paralvinella*, 15 to 26 in *P. Miralvinella* and 5 to 32 in *P. Nautalvinella*) (Desbruyères and Laubier, 1993; Jollivet and Hourdez, 2020; Han et al., 2021). Based on these diagnostic traits, Han et al. related the new indian species *P. mira* to the subgenus *Miralvinella* by considering the pointed shape of the sexual tentacles, the shape of gills and the presence of digitiform lobes on the notopodia. The authors still noted peculiarities, such as the first three setigerous segments, which are not fused in *P. mira*, and the insertion of numerous slender tentacles on the buccal apparatus (Han et al., 2021).

Our phylogenetic analyses however allow us to suggest a number of revisions to the taxonomy proposed by Desbruyères and Laubier within the genus *Paralvinella*.

Firstly, the subgenus *Nautalvinella* as described by Desbruyères & Laubier is also no longer monophyletic. If T9 (or T7) is kept as the most probable species trees, the species *P. p. irlandei* and *P. p. pandorae* are sister to the *Alvinella* and *Paralvinella* clades, and can be grouped in a new genus, possibly *Nautalvinella* as proposed by Stiller et al. (2020). Thus, the acquisition of comb-like gills, the absence of a pair of sexual tentacles on the buccal apparatus of males (which may be compensate by the building of a reproductive cocoon, D. Jollivet, pers. obs.) and the lack of digitiform notopodial lobes, which are the main elements in Desbruyères & Laubier’s diagnosis of the subgenus *Nautalvinella* (Desbruyères and Laubier, 1993) a symplesiomorphy.

Secondly, the phylogeny of the species which should belong to the subgenera *P. Miralvinella* and *P. Paralvinella* is well resolved. These species form a monophyletic clade in which *P. hessleri* and *P. mira* are positioned together with the *P. Paralvinella* species. The species *P. mira*, as suggested by Han et al. (2021), effectively groups with *P. hessleri*, which is the only other species attributed to *P. Miralvinella*. Unfortunately, molecular data are not available for the rarer species *P. dela* and *P. bactericola* to confirm whether they belong to the same grouping. According to Han et al., *P. mira* is morphologically closer to *P. hessleri* than to *P. dela* or *P. bactericola* when considering the shape of the buccal apparatus (stronger tentacles in *P. mira* and *P. hessleri*) and the position of the first uncinigerous neuropodial tori on chaetiger 16 or 18 in *P. mira* and *P. hessleri vs.* 32 for *P. dela* and *P. bactericola* (Han et al., 2021; Jollivet and Hourdez, 2020). The geographical distribution of the species also agrees with this view, as *P. mira* and *P. hessleri* are found in the Indian Ocean and south-west Pacific Ocean, while *P. dela* and *P. bactericola* are sister species inhabiting the Juan de Fuca Ridge and the Guaymas basin in the eastern Pacific Ocean (Desbruyères and Laubier, 1993; Jollivet and Hourdez, 2020). Based on our results, the subgenus *P. Paralvinella* is however no longer monophyletic since *P. mira* and *P. hessleri* are included in the middle of this group. This clearly indicates that the subgenus *P. Miralvinella* is no longer valid and that three-lobed tentacles of the mouth apparatus would then represent a symplesiomorphy, derived later as pointed tentacles in *P. mira*, *P. hessleri*, *P. dela* and *P. bactericola*.

Finally, if T9 represents the true species tree, then *P. unidentata* is closer to the two *Alvinella* species despite they do not share the same vent habitat, the former living in colder conditions more likely similar to those experienced by the *P. pandorae* worms. As a consequence, some specific traits such as the lack of sexual tentacles or the shape of the gills (comb-like gills) can be viewed as symplesiomorphies (possibly kept because of less constrained habitats) or may have arisen as a result of gene transfer between the *P. unidentata* and *P. pandorae* ancestor lineages.

### 4.3 Biogeographic history of alvinellid worms in and outside of the Pacific Ocean

Our molecular dating estimates rely solely on two calibration points. The first point corresponds to the recent opening of the Manus (3-4 Ma) and Lau (1-2 Ma) basins, which are assumed to open separately after the subduction of the Solomon Ridge and the fossilization of the South Fiji basin (Boulart et al., 2022). The second refers to the subduction of the Farallon Plate beneath the North American Plate, which caused a vicariant event between hydrothermal vent communities of the Eastern Pacific between 34 and 23 My (Tunnicliffe, 1988). This vicariant event is particularly interesting in the case of the Alvinellidae, since it separates three pairs of sibling species: *P. palmiformis* and *P. grasslei*, *P. p. irlandei* and *P. p. pandorae*, and *P. dela* and *P. bactericola* (Tunnicliffe, 1988; Desbruyères and Laubier, 1993; Jollivet and Hourdez, 2020). Most of the species of the three putative genera of the family Alvinellidae, namely *A. pompejana*, *A. caudata*, *P. p. irlandei*, *P. p. pandorae*, *P. grasslei*, *P. palmiformis*, *P. bactericola* and *P. dela* are endemic to the East Pacific Rise or the Juan de Fuca Ridge (Desbruyères and Laubier, 1986). The most parsimonious explanation for this high species richness regarding the phylogeny is an origin of the family in the Eastern Pacific Ocean. This is consistent with the biogeography of the sister family Ampharetidae, for which species diversity was the greatest in the eastern Pacific. This may suggest a common origin for both families in the now-extinct or subducted Pacific-Farallon and Mathematician ridges. These ridges have been active since the Early Jurassic (200 Ma) and have played a central role in the initial dispersal of the hydrothermal vent fauna in the Pacific (Mammerickx et al., 1980; Bachraty et al., 2009). The split between the Alvinellidae and Ampharetidae occurred between 110 and 162 Ma according to the CIR model, at a time when high spreading rates and off-ridge volcanism formed sub-aerial volcanic environments, which are now submerged in the Pacific (Röhl and Ogg, 1996). These emerged masses may have served as refuges for vent faunas during episodes of deep-sea anoxia during the Aptian to the Cenomanian ages (125 to 94 Ma) (Jacobs and Lindberg, 1998).

Depending on the model used for the molecular dating, the subsequent radiation of the family Alvinellidae occurred between 99 Ma (uncorrelated gamma model, 56-180 Ma) or 91 Ma (CIR model, 95% confidence interval 77-112 Ma). These estimates suggest that alvinellid worms have a history of speciation dating back to the Cretaceous period as previously proposed by Haymon et al. (1984); Vrijenhoek (2013). Using 278 orthologous genes or the *Cox1* mitochondrial gene, Jollivet and Hourdez proposed two dates for the radiation of the Alvinellidae, one at 198 Ma and the other at 72 Ma (Jollivet and Hourdez, 2020). Our estimates, although a little bit older, are in line with this second dating of Jollivet and Hourdez at 72 Ma, established from *Cox1* with calibration points on two pairs of sibling species. The radiation is therefore likely to follow the Cenomania/Turonian extinction phase, about 94 Ma. This major anoxic/dysoxic event has been argued to have created opportunities for modern taxa to invade deep environments (Thomas, 2007; Vrijenhoek, 2013) and are in agreement with vent fossil record exhibiting important transitions of the chemosynthetic faunas during the late Mesozoic (145-66 Ma) (Vrijenhoek, 2013).

However, the molecular dating of the tree’s root (radiation of the Terebelloformia, Fig. 5) appears much younger than the fossil records. We used a soft *prior* on the root’s age between 330 Ma +/- 200 My, in line with the abundance of terebellid fossils from the Carboniferous/Devonian era (Sepkoski Jr., 2002), but Thomas and Smith (1998), Little et al. (1999), and Vinn and Toom (2014) have proposed that ichnofossils dating back to Paleozoic (older than 440 Ma) could be attributed to the emergence of the Terebelliformia. There is therefore a discrepancy between the molecular dating done here (even using a older root’s age of 420 Ma +/- 100 My) and the fossil records. This should be addressed by including other fossil-dated outgroups, as recent speciation events used as calibration points could rejuvenate the molecular dating of the Alvinellidae radiation.

Since their radiation, the alvinellid worms expanded over the whole Pacific and the Indian Ocean (Han et al., 2021). The alternation in the phylogeny of eastern Pacific species (*A. pompejana*, *A. caudata*, *P. p. irlandei*, *P. grasslei*, *P. palmiformis*, *P. sulfincola*) and western Pacific (*P. unidentata*, *P. hessleri*, *P.* sp. nov., *P. fijiensis*) suggests however that the colonization of the Pacific Ocean must have occurred several times (Desbruyères and Laubier, 1993). There are indeed several dates of divergence between the western and eastern Pacific species ranging from 42 (*P. sulfincola vs. P. fijiensis*) to 87 Mya (*Alvinella vs. P. unidentata*). *P. dela* and *P. bactericola*, two eastern Pacific species missing in the phylogeny, are likely closer to *P. mira* and *P. hessleri* according to morphological characters. This implies that the oldest date of the western Pacific colonization appears to be between 65 and 35 My, which may correspond to the birth and spread of the Kula ridge (between 60 and 43 My (Smith, 2003)). This ridge has been indeed proposed to represent a bridge between the western and eastern Pacific hydrothermal basins, which are now isolated from each other (Hessler and Lonsdale, 1991; Bachraty et al., 2009). The second colonization of the Southwest Pacific region (leading to the separation of *P. fijiensis* and *P. sulfincola*) should have occurred between 45 and 22 My (54-18), potentially concurrent with the subduction of the Pacific plate under the Ontong-Java plateau between 45 and 30 Ma and the subsequent opening of modern-day back-arc basins (Schellart et al., 2006). The recent description of *P. mira* in the Indian Ocean, at Wocan and Daxi vents, shows that the alvinellid worms have also spread outside the Pacific Ocean. The divergence between *P. mira* and *P. hessleri* is dated back to 35 My (27-48) and could be an argument for the recent colonization of the Indian Ocean. The connection between the western Pacific and the Indian Ocean through Northern Australia was closed between 60 and 40 Ma, with the advance of the Philippines and the opening of the China Sea (Parker and Gealey, 1985; Moalic et al., 2012). Although the tectonic history of this region makes it difficult to reconstruct ridge connectivity between oceanic basins, the colonization of the Indian Ocean by Alvinellidae was much more likely made possible through the Antarctic Ridge, whose spread started about 50 Ma and connects the Indian Ocean with the Pacific Ocean (Parker and Gealey, 1985). This ridge is indeed a link to most of the vent sites, from the EPR to the mid-Atlantic Ridge through the Indian Ocean (Moalic et al., 2012).

### 4.4 Unusual compositional amino-acid biases for P. p. irlandei

Phylogenetic reconstruction is sensitive to differences in species sequence composition, which can lead in particular to long-branch attraction. Alvinellidae colonize contrasting thermal environments, which is known to have an effect on the amino-acid composition of proteins between mesophilic and thermophilic eukaryotes (Wang and Lercher, 2010; van Noort et al., 2013; Bock et al., 2014).

Although almost no bias in the codon use between alvinellid worms Fontanillas et al. (2017), the evaluation of the amino-acid composition bias in the alvinellid orthologous gene clusters shows strong disparities between species (see supplementary data Fig. 7). In particular, *P. p. irlandei* shows a very different amino-acid composition when compared to other species (alvinellid species but also the outgroup species), with great differences in indices such as CvP bias, Serine bias and purine load: criteria which have been suggested to discriminate between warm and cold species (Hickey and Singer, 2004; Fontanillas et al., 2017). Therefore, we carefully filtered out genes that exhibited a high compositional heterogeneity in our phylogenetic analyses. We also ensured that each gene alignment contained a sequence of *P. p. irlandei* in the analysis, although most of the *P. p. irlandei* sequences were on average shorter and more fragmented (158,783 total nucleotide sites and 292,610 amino acid sites). This limits the available phylogenetic information for this species compared to other alvinellid species of the dataset, and may have altered the placement of this species in the alvinellid tree. One might indeed expect that the strong compositional bias found in *P. p. irlandei* would tend to make this species diverge more rapidly among the Alvinellidae in the phylogenetic reconstruction. The positioning of *P. p. irlandei* in the alvinellid tree could be highly gene-dependent when estimating tree topologies (and to a lesser extent when fitting constrained gene trees). One must therefore be cautious with the good amino-acid scores of the T7, T8 and T9 topologies, which place *P. p. irlandei* sister to other Alvinellidae species, as it may be a typical consequence of long-branch attraction, and needs to compare these scores with those of the nucleic acids.

### 4.5 Phylogenomic incongruence reveals a complex evolutionary history of Alvinellidae

Rapid radiation, as displayed by the Alvinellidae, can lead to a high rate of incomplete lineage sorting (ILS) in the early steps of speciation and results in larger gene tree estimation errors (GTEE) (Molloy and Warnow, 2018). In this case, coalescent-based methods, which account for different gene histories, should give better results than concatenation methods for estimating the true species tree. However, the superiority of coalescent methods depends on the confidence level put in gene tree estimation. For low or medium ILS or inter-species gene flow, the concatenation method, which benefits from longer alignments, allows the reduction of GTEE, and is still more reliably in practice (Roch and Warnow, 2015; Molloy and Warnow, 2018). In the case of high ILS or large amounts of inter-species gene flows, coalescent methods are preferable provided that GTEE remains low, particularly with the longest possible gene alignments (Nute et al., 2018).

As shown in figure 3b. and 3d., considering genes with the highest phylogenetic signal, and thus the lowest GTEE, did not allow to exclude some gene tree topologies compared to genes with a lower phylogenetic signal. The two similar distributions indicate that GTEE remained rather low due to our strategy of evaluating constrained topologies. ILS is therefore likely to be mainly responsible for the distribution of gene trees. However, the distribution of tree topologies is not random with T6 and T9 being more often supported. From this gene distribution, two alternative scenarios where *P. unidentata* is either closer to the *Alvinella* species (mainly supported by T9), or closer to *P. p. irlandei* (mainly supported by T6) seem to be equiprobable, as shown in figure 3a. and 3c. Such a bimodality is expected if one of these topologies is reflecting the true species tree, while the other is the consequence of specific inter-species gene flow between ancestors. Yet these two scenarios are not directly compatible with one another if we consider that T9 is the true species tree. Nevertheless, the hypothesis of a massive gene introgression between these two ancestors may explain why the T7 topology is favored in the coalescent framework using constrained gene trees resolution (Figure 4). T7, which groups *P. unidentata* with other *Paralvinella*, is an intermediate topology between T6 and T9 that could easily explain the different gene histories. Some of them would be linked to the species formation, and some others with secondary asymmetric gene introgression from either the ancestor of *P. unidentata* to the ancestor of *P. p. irlandei* (resulting in T6), or between ancestors of *P. unidentata* and the *Alvinella* lineage (resulting in T9). As such, it could reconcile the main phylogenetic relationships that appear in the multi-gene analysis. Interestingly, T7 is also one of the most supported topology among amino-acid encoded genes (Figure 3d.), despite not being statistically over-represented compared to a random distribution (p-value=77%).

Although we propose T9 as the main species phylogeny, our results show that ancestral polymorphism and high introgression between species has blurred the phylogenetic signal during the alvinellid radiation. The sequencing effort, which includes several hundred genes, is already very substantial and it is unlikely that increasing the number of sequences will improve this result significantly. In the future, mapping the genes carrying different phylogenetic signals onto an alvinellid genome could enable us to see which regions of the genome are associated with specific evolutionary histories. This question should be resolved before considering to revise the genus name of *P. p. irlandei*, *P. p. pandorae* and *P. unidentata*.

## 5. Conclusion

To conclude, our phylogenetic analysis of the alvinellid worms led support the polyphyly of the *Paralvinella* genus (T9 topology), with *P. p. irlandei* being sister to other alvinellid species). As such, both *P. pandorae pandorae* and *P. pandorae irlandei* should belong to a new genus (possibly *Nautalvinella*). This fits well with the more simplistic morphology of these worms and their opportunistic strategy to colonize the vent environment as they represent ’pioneer species’. The positioning of *P. unidentata* is more uncertain as it may group with the *Alvinella* species. Such uncertainty is mostly due to the fact that most of genes have different phylogenetic histories, which is probably a consequence of the rapid radiation of several lineages from an ancestral worm population, resulting in high incomplete lineage sorting and intraspecific gene introgression. As a consequence, two alternative topologies (T6 and T7) are also well supported, and in agreement with some morphological traits depending on whether some of them represent synapomorphies or symplesiomorphies. The phylogenetic positioning of the other *Paralvinella* species is robust and rules out the idea that *P. hessleri* and *P. mira* can be grouped in the subgenus called *Miralvinella*. As a consequence, the gill shape of alvinellid worms seems to represent a good proxy to separate species into different genera. However, this trait is likely the result of natural selection in the face of local hypoxia, and together with the amino-acid residue biases found in proteins are likely to have emerged very early in their evolution as an adaptation to contrasting vent conditions.

## 6. Credit authorship contribution statement

Pierre-Guillaume Brun - Data Curation, Methodology, Formal Analysis, Writing original draft and review. Stéphane Hourdez - Funding acquisition, Resources, Reviewing original draft. Marion Ballenghien - Methodology. Yadong Zhou - Resources, Reviewing original draft. Jean Mary - Supervision, Funding acquisition, Reviewing original draft. Didier Jollivet - Supervision, Project administration, Funding acquisition, Resources, Writing original draft and review.

## 7. Supplementary Material

Data available from the Dryad Digital Repository: https://doi.org/10.5061/dryad.dbrv15f6f.

