## supplementary data for "A step in the deep evolution of Alvinellidae (Annelida: Polychaeta): A phylogenomic comparative approach based on transcriptomes"

| | Raw sequencing reads ($x10^6$) and bases ($x10^9$) | Sequencing reads ($x10^6$) and bases ($x10^9$) after filtration | Assembled transcripts ($x10^3$) | % prokaryotic contamination | ORF ($x10^3$) | Busco % complete or % fragmented transcripts | Orthologous genes |
|---|---|---|---|---|---|---|---|
| *Pectinaria gouldii* | / (1) | / (1) | 55 | 0.31 | 40 | 53.5/21.6 | 1235 |
| *Melina palmata* | 334.5/50.51 | 223.8/29.4 | 872 | 0.07 | 515 | 99.0/0.4 | 1939 |
| *Terebellidae. sp.* (2 individuals) | 2014.3/304.2 | 1545.1/195.4 | 2340 | 0.08 | 1308 | 95.0/4.0 | 1879 |
| *Neoamphitrite edwardsii* | 280.9/42.4 | 186.6/24.4 | 366 | 0.08 | 183 | 98.1/0.3 | 1908 |
| *Anobothrus* spp. | 76.8/7.7 | 56.6/5.4 | 432 | 0.08 | 193 | 90.3/6.3 | 1753 |
| *Amphisamytha carldarei* | 78.5/7.9 | 43.8/4.2 | 280 | 0.07 | 126 | 76.4/19.7 | 1670 |
| *Amphicteis gunneri* | 95.9/9.6 | 71.3/6.9 | 408 | 0.07 | 182 | 86.1/9.9 | 1688 |
| *Hypania invalida* | 49.5/4.7 | 27.1/2.5 | 195 | 0.06 | 124 | 97.4/1.8 | 1905 |
| *Alvinella caudata* | 230.1/34.8 | 154.9/20.4 | 394 | 0.10 | 299 | 99.2/0.3 | 1925 |
| *Alvinella pompejana* | / (2) | / (2) | 45 | 6.52 | 39 | 94.9/2.8 | 1871 |
| *Paralvinella pandorae irlandei* | 13.8/1.0 | 1.0/0.7 | 195 | 0.27 | 80 | 35.4/31.1 | 1009 |
| *Paralvinella unidentata* 1 | 311.1/31.1 | 244.3/23.6 | 488 | 0.50 | 369 | 98.5/0.6 | 1942 |
| *Paralvinella unidentata* 2 | 407.2/61.5 | 321.5/41.3 | 345 | 0.22 | 227 | 96.1/2.6 | 1907 |
| *Paralvinella unidentata* 3 | 316.9/47.9 | 248.8/31.8 | 374 | 0.27 | 488 | 98.3/0.8 | 1925 |
| *Paralvinella palmiformis* | 270.6/27.1 | 221.7/21.3 | 372 | 0.30 | 194 | 98.7/0.4 | 1916 |
| *Paralvinella grasslei* (2 individuals) | 56.8/5.5 | 46.1/4.4 | 223 | 0.23 | 132 | 76.2/19.1 | 1595 |
| *Paralvinella mira* | 13.4/2.0 | 11.0/1.5 | 127 | 0.16 | 86 | 91.0/4.5 | 1849 |
| *Paralvinella hessleri* 1 | 418.6/41.9 | 330.5/31.9 | 467 | 0.29 | 268 | 97.8/1.3 | 1925 |
| *Paralvinella hessleri* 2 | 685.2/103.5 | 531.0/68.3 | 331 | 0.22 | 179 | 96.5/2.4 | 1890 |
| *Paralvinella sulfincola* | / (1) | / (1) | 21 | 0.41 | 32 | 68.1/14.6 | 1126 |
| *Paralvinella sp.* nov. (2 individuals) | 894.7/135.1 | 669.4/85.4 | 560 | 0.35 | 276 | 83.4/13.3 | 1460 |
| *Paralvinella fijiensis* Manus Basin | 1092.2/164.9 | 858.9/109.8 | 432 | 0.34 | 269 | 98.5/0.6 | 1921 |
| *Paralvinella fijiensis* 1 Lau Basin | 326.5/32.6 | 252.8/24.4 | 496 | 0.52 | 359 | 98.6/0.5 | 1938 |

| | Raw sequencing reads ($x10^6$) and bases ($x10^9$) | Sequencing reads ($x10^6$) and bases ($x10^9$) after filtration | Assembled transcripts ($x10^3$) | % prokaryotic contamination | ORF ($x10^3$) | Busco % complete or % fragmented transcripts | Orthologous genes |
|---|---|---|---|---|---|---|---|
| *Paralvinella fijiensis* 2 Lau Basin | 493.0/74.4 | 384.1/49.4 | 347 | 0.24 | 223 | 97.3/1.9 | 1866 |
| *Paralvinella fijiensis* 3 Lau Basin | 430.3/65.0 | 336.1/43.3 | 325 | 0.24 | 198 | 97.4/1.5 | 1892 |

**Table 1. *Metrics of the different assembly steps and identification of orthologous genes.*** Reads are filtered with Fastp. Transcript assembly is performed with Trinity. Prokaryotic transcript cleaning is performed with Kraken. ORF delineation and trimming of 5' and 3' UTRs is performed with Transdecoder. Transcriptome completeness assessment is performed with Busco in comparison with the ODB9 Metazoa database (978 Buscos). Orthologous gene clusters are retrieved with Orthograph, based on a database of 1997 orthogroup genes identified in lophotrocozoa as single-copy genes. (1) Transcriptomes provided by Didier Jollivet. (2) Transcripts retrieved with Augustus from the *A. pompejana* genome.

### Command lines

#### FASTP

```
fastp -i $1 -o reads_fastp_R1.fastq.gz -I $2 -O reads_fastp_R2.fastq.gz $phred
--compression=9 --detect_adapter_for_pe --adapter_fasta adapters-primers.txt -n
10 -q 25 -u 50 -l 50 -y 10 --cut_right --cut_right_window_size=5 --
cut_right_mean_quality 30 --correction --overlap_len_require 20 --
overlap_diff_limit 2 --overlap_diff_percent_limit 5 --cut_front --
cut_front_window_size 1 --cut_front_mean_quality 25 --trim_poly_g --
poly_g_min_len 20 --trim_poly_x --poly_x_min_len 20
```

#### TRINITY ASSEMBLY

```
Trinity --seqType fq --max_memory 130G --left reads_fastp_kraken_R1.fq.gz --
right reads_fastp_kraken_R2.fq.gz --CPU 8 --min_contig_length 50 --output
transcriptome_trinity --full_cleanup
```

#### CAP3

```
cap3 transcriptome.fasta -o 50 -p 99 > transcriptome.cap3.fasta
```

| | Number of nucleotide sites (% of MSA length) | Number of amino acid sites (% of MSA length) | Number of genes in nucleotide orthologous groups (%) | Number of genes in amino-acid orthologous groups (%) |
|---|---|---|---|---|
| *Pectinaria gouldii* | 245,846 (49) | 139,067 (50) | 474 (72) | 515 (74) |
| *Melina palmata* | 449,432 (90) | 251,511 (91) | 644 (98) | 694 (99) |
| *Terebellidae. sp.* (2 individuals) | 442,088 (89) | 247,729 (89) | 639 (97) | 688 (98) |
| *Neoamphitrite edwardsii* | 433,188 (87) | 251,135 (90) | 623 (95) | 688 (98) |
| *Anobothrus* spp. | 422,984 (85) | 234,701 (84) | 630 (96) | 679 (97) |
| *Amphisamytha carldarei* | 395,516 (79) | 215,149 (77) | 633 (97) | 671 (96) |
| *Amphicteis gunneri* | 424,408 (85) | 234,900 (85) | 633 (97) | 678 (97) |
| *Hypania invalida* | 441,160 (88) | 247,978 (89) | 635 (97) | 687 (98) |
| *Alvinella caudata* | 461,042 (92) | 255,557 (92) | 657 (100) | 699 (100) |
| *Alvinella pompejana* | 432,660 (87) | 237,045 (85) | 645 (98) | 687 (98) |
| *Paralvinella pandorae irlandei* | 292,610 (59) | 158,783 (57) | 657 (100) | 699 (100) |
| *Paralvinella unidentata* 1 | 448,586 (90) | 248,039 (89) | 657 (100) | 697 (100) |
| *Paralvinella unidentata* 2 | 451,370 (90) | 247,166 (89) | 652 (99) | 695 (99) |
| *Paralvinella unidentata* 3 | 445,690 (89) | 250,479 (90) | 654 (100) | 694 (99) |
| *Paralvinella palmiformis* | 445,686 (89) | 245,611 (88) | 652 (99) | 693 (99) |
| *Paralvinella grasslei* (2 individuals) | 356,258 (71) | 198,753 (72) | 556 (85) | 592 (85) |
| *Paralvinella mira* | 428,682 (86) | 236,672 (85) | 646 (98) | 688 (98) |
| *Paralvinella hessleri* 1 | 452,758 (91) | 250,575 (90) | 652 (99) | 692 (99) |
| *Paralvinella hessleri* 2 | 448,750 (90) | 245,678 (88) | 649 (99) | 688 (98) |
| *Paralvinella sulfincola* | 290,084 (58) | 155,946 (56) | 540 (82) | 573 (82) |
| *Paralvinella sp.* nov. (2 individuals) | 346,800 (69) | 188,942 (68) | 558 (85) | 594 (85) |
| *Paralvinella fijiensis* Manus Basin | 454,852 (91) | 251,483 (90) | 655 (100) | 697 (100) |
| *Paralvinella fijiensis* 1 Lau Basin | 457,354 (92) | 253,438 (91) | 657 (100) | 698 (100) |
| *Paralvinella fijiensis* 2 Lau Basin | 455,898 (91) | 249,592 (90) | 656 (100) | 697 (100) |
| *Paralvinella fijiensis* 3 Lau Basin | 452,402 (91) | 249,298 (90) | 651 (99) | 694 (99) |

**Table 2. *Number of sites/genes for each transcriptome in the concatenated supermatrices or gene trees.***

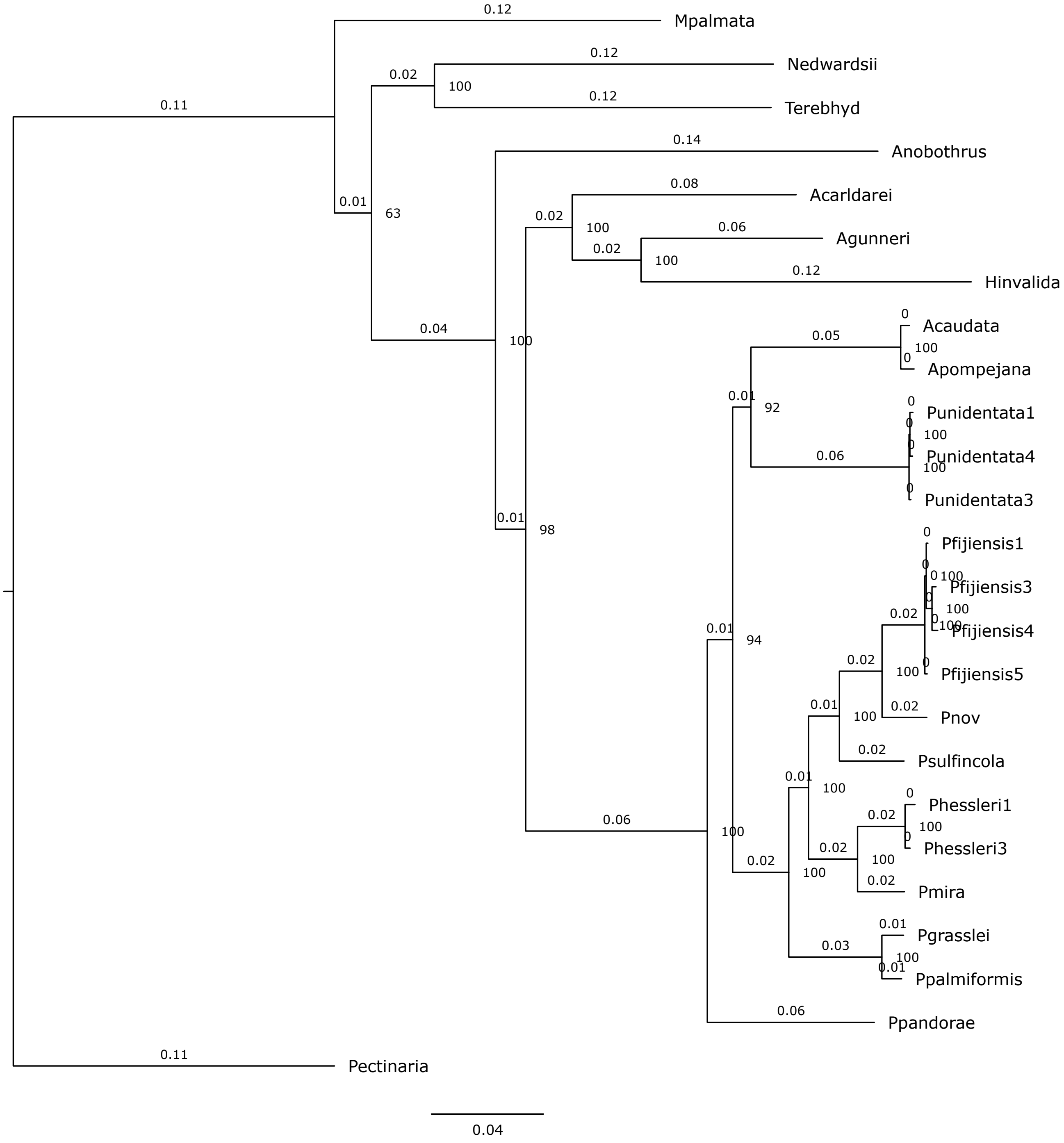


**Figure 1.** ***Phylogeny obtained from 657 concatenated nucleotide genes*** (499,036 sites) shared by at least 20 transcriptomes. IQ-TREE 2.0.3, partitioned model. 1000 ultrafast bootstraps.

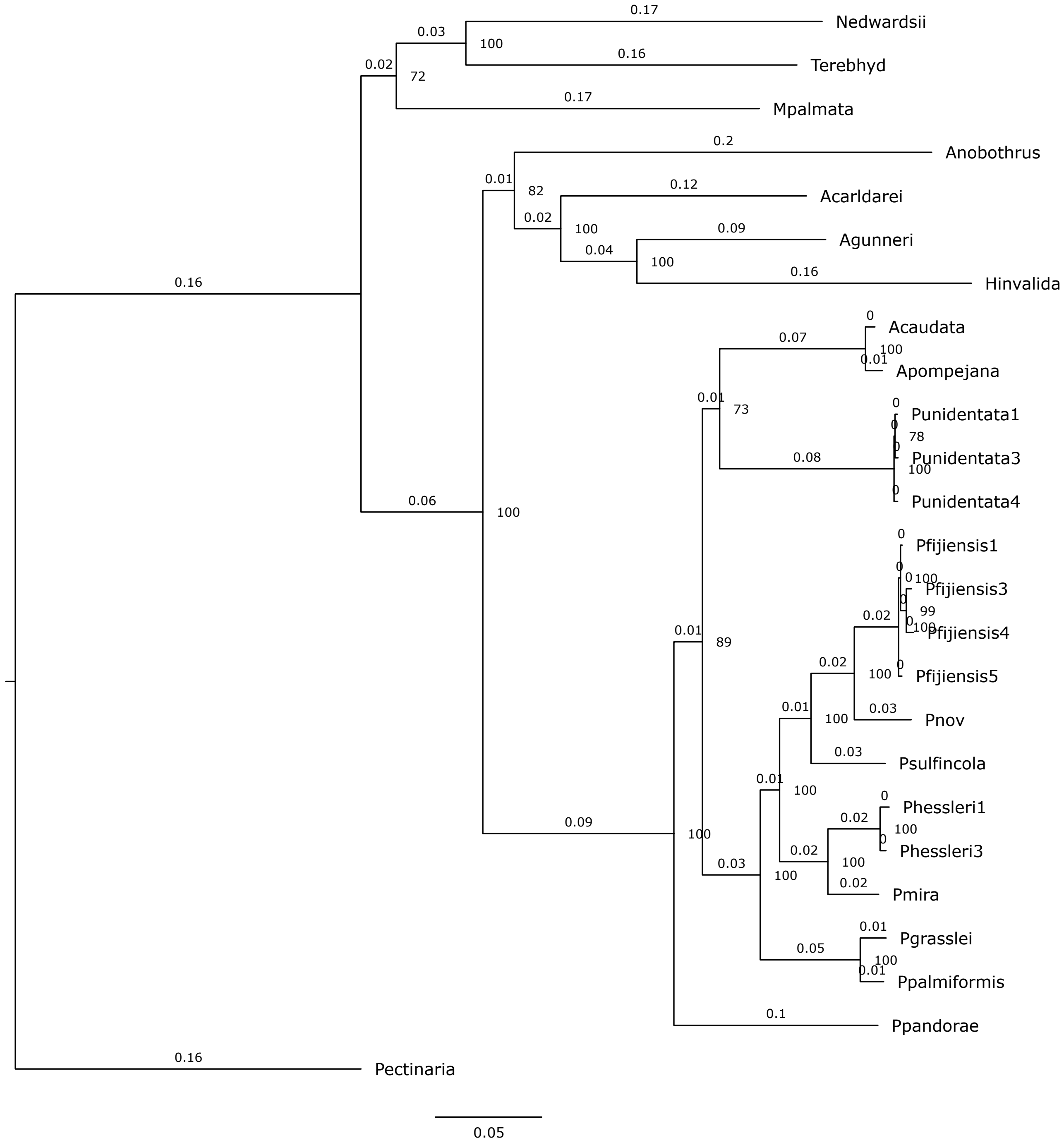


**Figure 2.** ***Phylogeny obtained from 699 concatenated amino acid genes*** (277,900 sites) shared by at least 20 transcriptomes. IQ-TREE 2.0.3, partitioned model. 1000 ultrafast bootstraps.

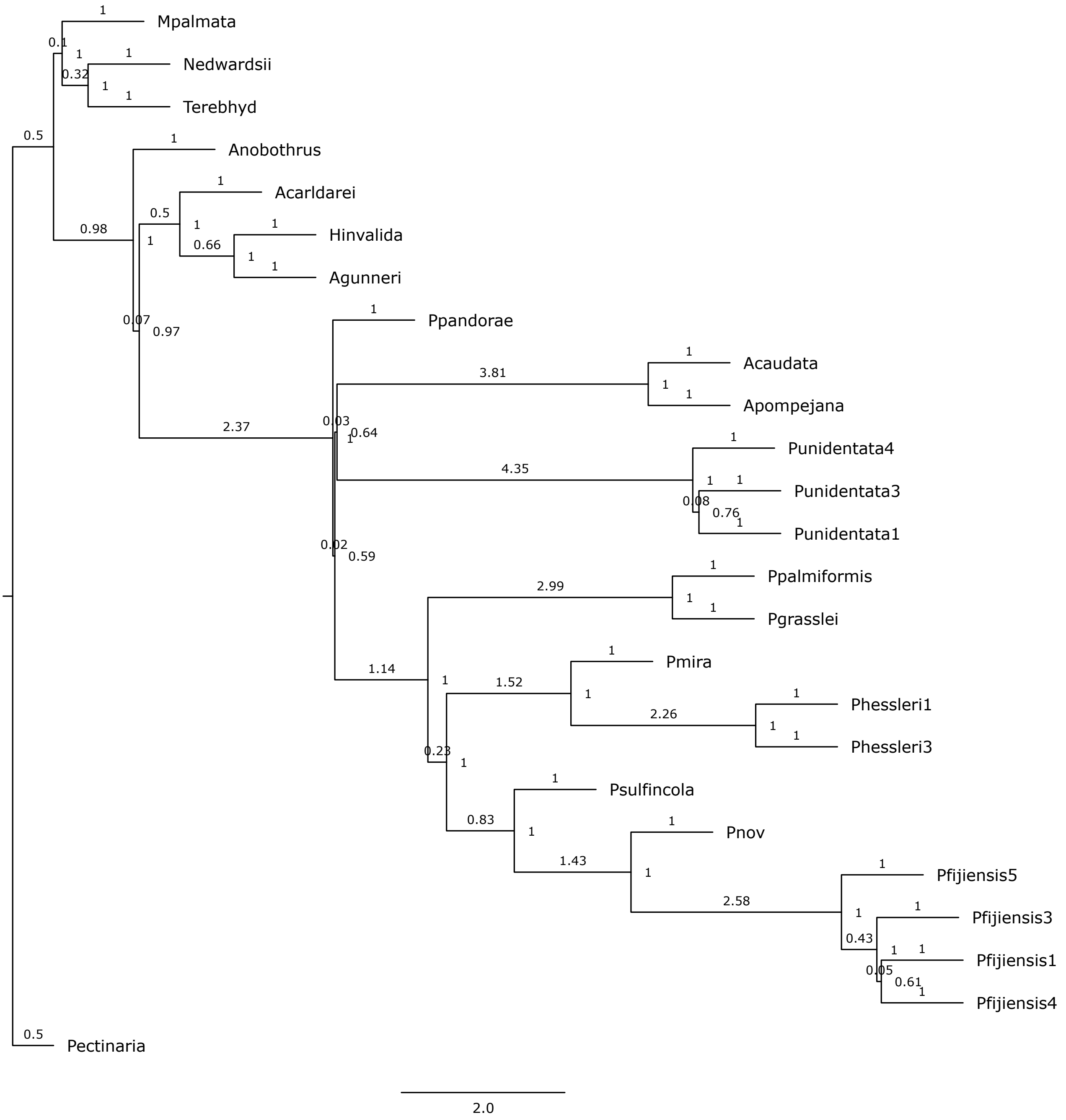


**Figure 3.** ***Phylogeny obtained from 657 nucleotide gene trees*** shared by at least 20 transcriptomes. Gene trees are obtained with IQ-TREE 2.0.3. The species tree is obtained with ASTRAL 5.7.8. Posterior probabilities are indicated at each node.

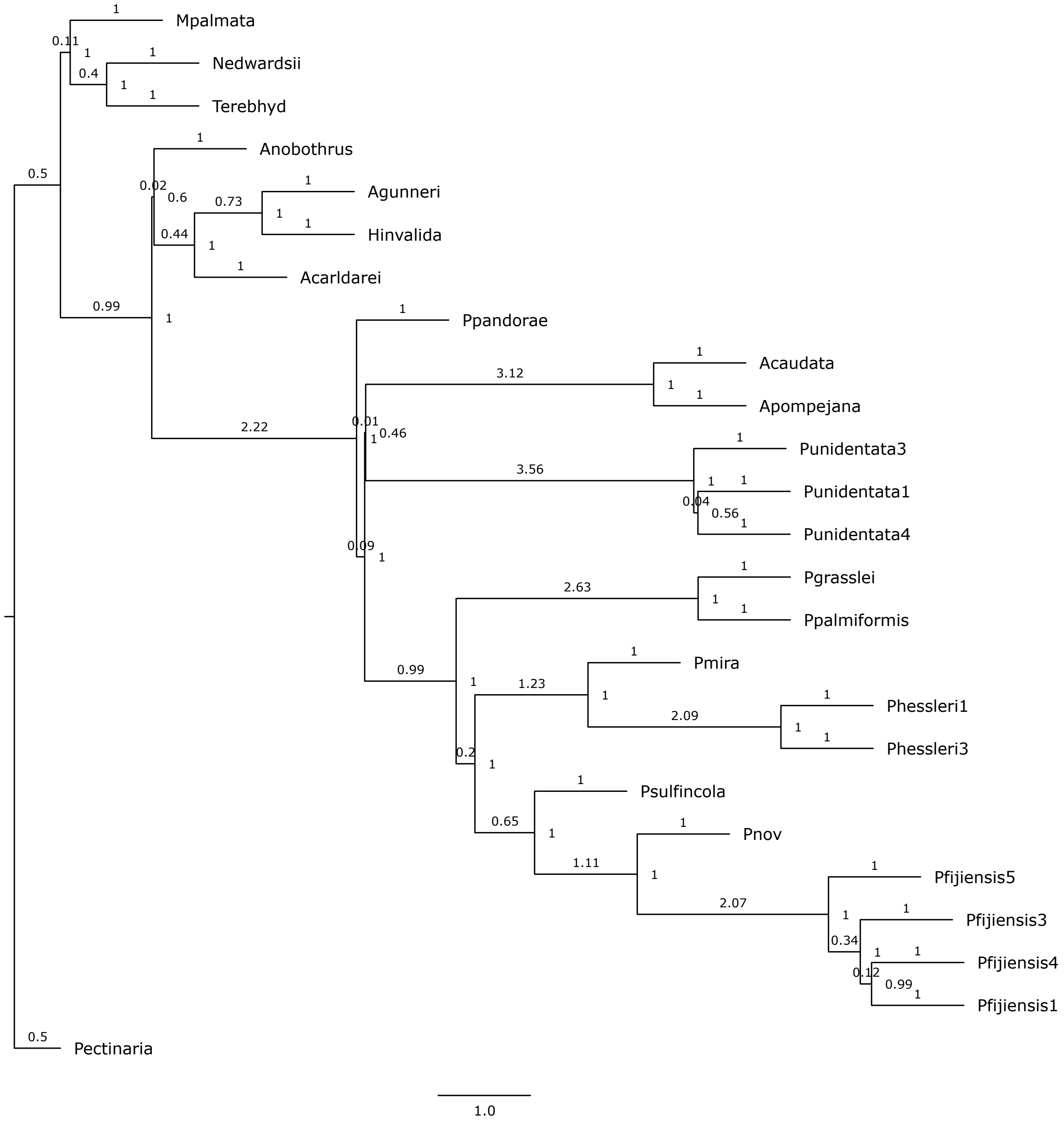


**Figure 4. *Phylogeny obtained from 699 amino acid coded gene trees*** shared by at least 20 transcriptomes. Gene trees are obtained with IQ-TREE 2.0.3. The species tree is obtained with ASTRAL 5.7.8. Posterior probabilities are indicated at each node.

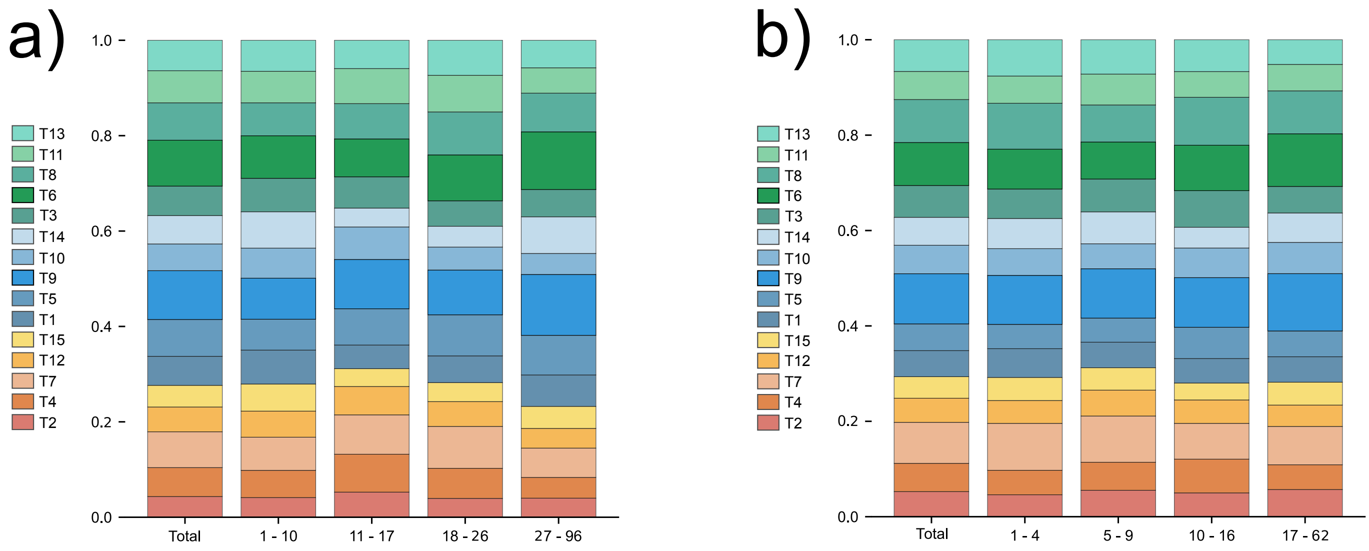


**Figure 5.** ***Topology weight across genes.*** (a) Distribution of genes' probabilities according to the number of parsimony-informative sites for nucleotide sequences. (b) Distribution of genes' probabilities according to the number of parsimony-informative sites for amino-acid sequences.

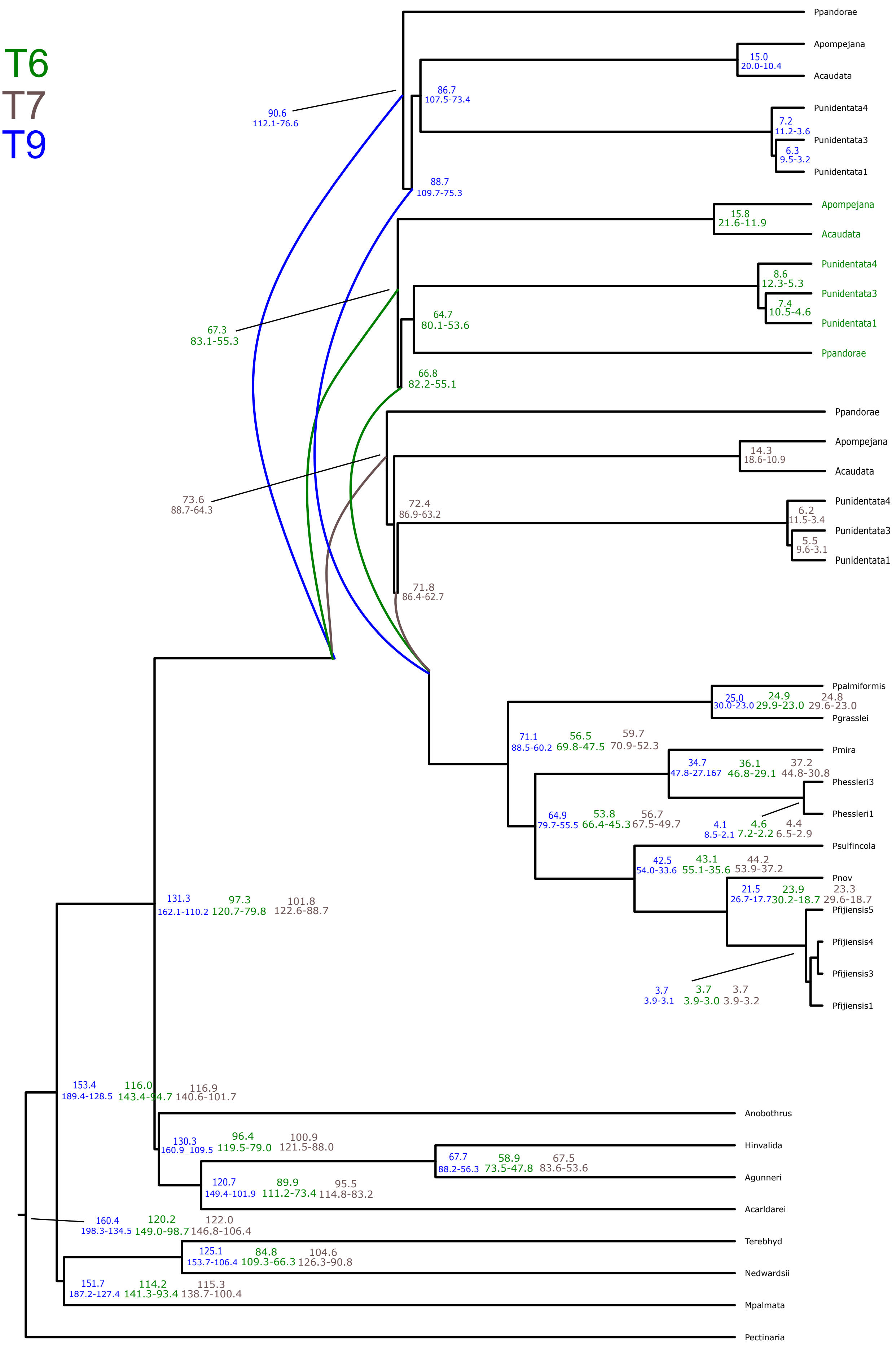


**Figure 6.** ***Age estimates according to the CIR model***, obtained with Phylobayes 4.1 under the three topologies T6, T7 and T9 with 95% confidence intervals. Date estimates for other molecular clock models (log Normal and Uncorrelated gamma) are given in supplementary material as chronograms.

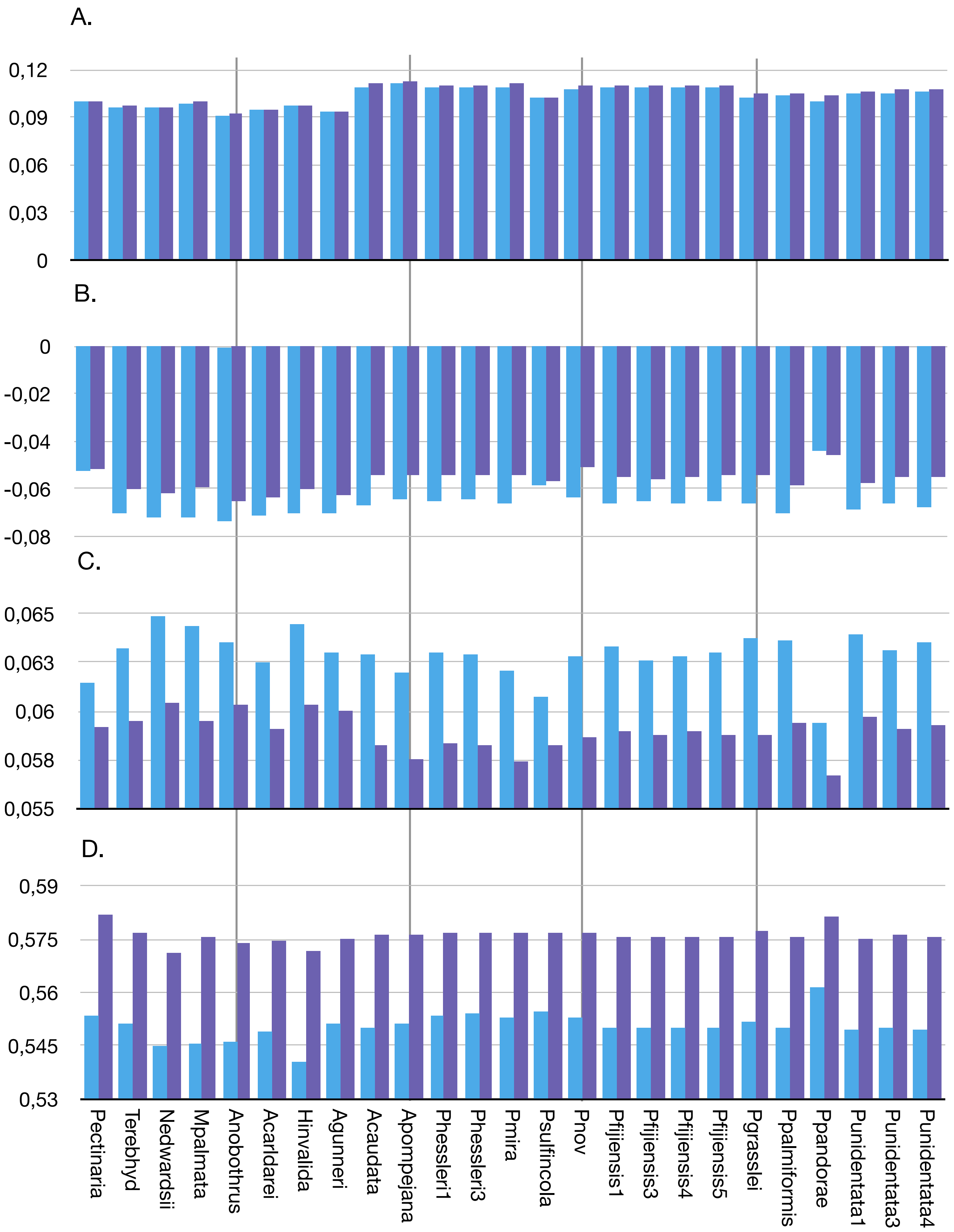


**Figure 7.** ***Composition of the transcriptomes.*** The blue bars correspond to the raw compositions of the total assembled transcripts, trimmed of the 5' and 3' UTRs and assigned to an ortholog group by Orthograph. The purple bars correspond to the compositions after filtration (composition/stationarity test as well as 3rd codon nucleotide removal for nucleotide-encoded genes). The sequences used for phylogenetic inference correspond to the filtered genes. **A.** PAYLE vs. DGMS bias. **B.** CvP bias (EDKR vs. GHNPQST). **C.** Serine bias. **D.** Purine load (AG vs. TC).
